# Functional Dissection of the Zdhhc5-GOLGA7 Protein Palmitoylation Complex

**DOI:** 10.1101/2025.04.28.651057

**Authors:** Martha A. Kahlson, Joan Ritho, João Victor Gomes, Haoqing Wang, Joel A. Butterwick, Scott J. Dixon

**Affiliations:** Department of Biology, Stanford University, Stanford, CA 94305, USA; Department of Pharmacology, Yale University, New Haven, CT 06510 USA; Stanford Cryo-EM Microscopy Center, Stanford University, Stanford, CA 94305, USA; Cryoelectron Microscopy, Nucleus at Sarafan ChEM-H, Stanford University, Stanford, CA 94305, USA

**Keywords:** S-acylation, necrosis, CIL56, cell death

## Abstract

Small molecules serve as valuable tools for probing non-apoptotic cell death mechanisms. The small molecule CIL56 induces a unique form of non-apoptotic cancer cell death that is promoted by a complex formed between zinc finger DHHC-type palmitoyltransferase 5 (ZDHHC5) and an accessory protein, GOLGA7. However, the structure, function, and regulatory role of this complex in cell death remain poorly understood. In this study, we employ biochemical purification, cryogenic electron microscopy (cryo-EM), homology modeling, mutagenesis, and functional assays to elucidate the structure of the Zdhhc5-GOLGA7 complex. We identify key conserved residues in both Zdhhc5 and GOLGA7 that are necessary for complex formation and to promote non-apoptotic cancer cell death in response to CIL56. These results provide new insights into the structure and function of a death-promoting protein complex.

## INTRODUCTION

Drugs that can induce cell death may be useful for the treatment of cancer. Synthetic small molecules that can induce cancer cell death in novel ways are therefore highly valued. A small molecule phenotype screen identified caspase independent lethal 56 (CIL56) as a novel inducer of non-apoptotic cancer cell death (1). CIL56 alters the morphology of multiple organelles, including the Golgi apparatus, the endoplasmic reticulum and the mitochondria, and is associated with perturbation of protein anterograde and retrograde protein trafficking (2). CIL56-induced cell death is prevented by 5-tetradecyloxy-2-furoic acid (TOFA), a small molecule inhibitor of de novo lipid synthesis (3). This implied a link between lipid metabolism and CIL56-induced cell death. Chemical genetic screening identified zinc finger DHHC-type palmitoyltransferase 5 (ZDHHC5) as an enzyme necessary for CIL56-induced cell death (2). ZDHHC5 is one of 23 mammalian enzymes that catalyze the *S*-acylation (“palmitoylation”) of hundreds of proteins in the cell, regulating protein localization, stability, and function (4, 5). Protein palmitoylation can regulate the execution of apoptosis and various forms of non-apoptotic cell death (6–8). How ZDHHC proteins regulate cell death is poorly understood.

The family of 23 mammalian ZDHHC enzymes are defined by an Asp-His-His-Cys (DHHC) active site motif. The terminal cysteine residue of this motif is necessary for lipid transfer to substrate proteins (9). Most ZDHHC-family enzymes localize to the endoplasmic reticulum or Golgi apparatus. However, ZDHHC5 is localized to the plasma membrane (PM) and ZDHHC8, ZDHHC20, and ZDHHC21 show partial PM localization as well (10–12). Some ZDHHC enzymes require accessory proteins to function. This was first established in *S. cerevisiae*, where the ZDHHC9 ortholog Erf2 requires the accessory protein Erf4 to function properly (13, 14). In mammalian cells, ZDHHC6 complexes with SELENOK, ZDHHC9 and ZDHHC5 complex with GOLGA7 (also known as GCP16), and ZDHHC5 also complexes with a GOLGA7 paralog, GOLGA7B (2, 14–18). These accessory proteins assist in substrate selection, stabilize enzyme intermediates, and enhance stability of their respective ZDHHC partners (13, 15, 18–20). The full spectrum of ZDHHC complexes that can form in mammalian cells and how these complexes regulate cell phenotypes is unclear.

We previously identified the ZDHHC5-GOLGA7 complex and showed that it was essential for CIL56-induced cell death (2, 3). In *ZDHHC5* gene-disrupted HT-1080 fibrosarcoma cells (“*ZDHHC5^KO^*”), re-expression of wild-type mouse Zdhhc5 (which shares 98% sequence identity with the human protein, **Supplementary Fig. S1*A***) fully restored sensitivity to CIL56. A Zdhhc5 enzyme-dead Cys to Ser active site mutant was able to interact with endogenous GOLGA7 but unable to restore cell death, implying that protein palmitoylation was essential for cell death. A Zdhhc5 mutant lacking three C-terminal cysteine residues was mis-localized inside the cell, unable to interact with endogenous GOLGA7, and unable to restore CIL56 sensitivity when expressed in *ZDHHC5^KO^* cells (2). How ZDHHC5 and GOLGA7 interact at the biochemical and structural levels to promote this cell death remains poorly understood.

Here, we use co-immunoprecipitation, functional assays, and cryogenic electron microscopy (cryo-EM) combined to map key residues necessary for the Zdhhc5-GOLGA7 complex to form and promote cell death in response to CIL56. We identify an “RNYR” sequence motif adjacent to the DHHC active site in Zdhhc5 and other ZDHHC-family enzymes as essential for interaction with GOLGA7. We also identify an “RDYS” motif found on GOLGA7 and GOLGA7B that is necessary for complex formation with ZDHHC5. Structural modeling by cryo-EM and AlphaFold 3 of the ZDHHC5-GOLGA7 complex reveals that the ZDHHC5^RNYR^ and GOLGA7^RDYS^ motifs contribute to the key interaction surfaces between these proteins. These results underscore structural elements necessary for ZDHHC5-GOLGA7 complex formation and the induction of non-apoptotic cell death.

## RESULTS

### GOLGA7 interacts with multiple Zdhhc family proteins

GOLGA7 was initially identified in complex with ZDHHC9 (16). As part of our studies, we showed using co-immunoprecipitation (co-IP) in 293T cells that endogenous GOLGA7 can complex not only with exogenous Zdhhc5 but also with the closely-related family member Zdhhc8 (2). The observation that GOLGA7 binds at least three mammalian ZDHHC enzymes initially prompted us to examine whether GOLGA7 could interact with additional ZDHHC family proteins, and to examine whether the information encoded in these interactions could be used to elucidate key protein features necessary for complex formation.

To explore the scope of GOLGA7 interactors, we overexpressed HA-tagged versions of all 23 mouse Zdhhc-family proteins (21) in 293T cells and assessed their ability to co-IP endogenous human GOLGA7 (**Fig. 1*A***). The decision to use mouse Zdhhc proteins was driven by the availability of this library of constructs spanning the entire family, and the fact that mouse Zdhhc proteins share on average 90% sequence identity with their human orthologs (**Supplementary Fig. S1*A***). These overexpression experiments were conducted in both established (2) 293T *ZDHHC5* gene-disrupted (knockout, “KO”) cell lines as well as in unmodified Control cells, hypothesizing that *ZDHHC5^KO^*cells might increase the availability of GOLGA7 for interaction with other overexpressed Zdhhc proteins. However, comparable results were obtained in both *ZDHHC5^KO^* and Control cells. Multiple independent pulldown experiments were performed for each HA-Zdhhc protein, using HA-Zdhhc5 as a positive control. Immunoblot analysis demonstrated substantial expression of all Zdhhc proteins, except for Zdhhc6 and Zdhhc12, which exhibited poor expression (**Supplementary Fig. S1*B*)**. The relative expression of each PAT was calculated, and densitometry quantification of GOLGA7 binding to each overexpressed Zdhhc protein, normalized to the Zdhhc5-GOLGA7 interaction (positive control), was summarized as a heatmap (**Fig. 1, *B* and *C***, **Supplementary Fig. S1*B***).

**Figure 1.**
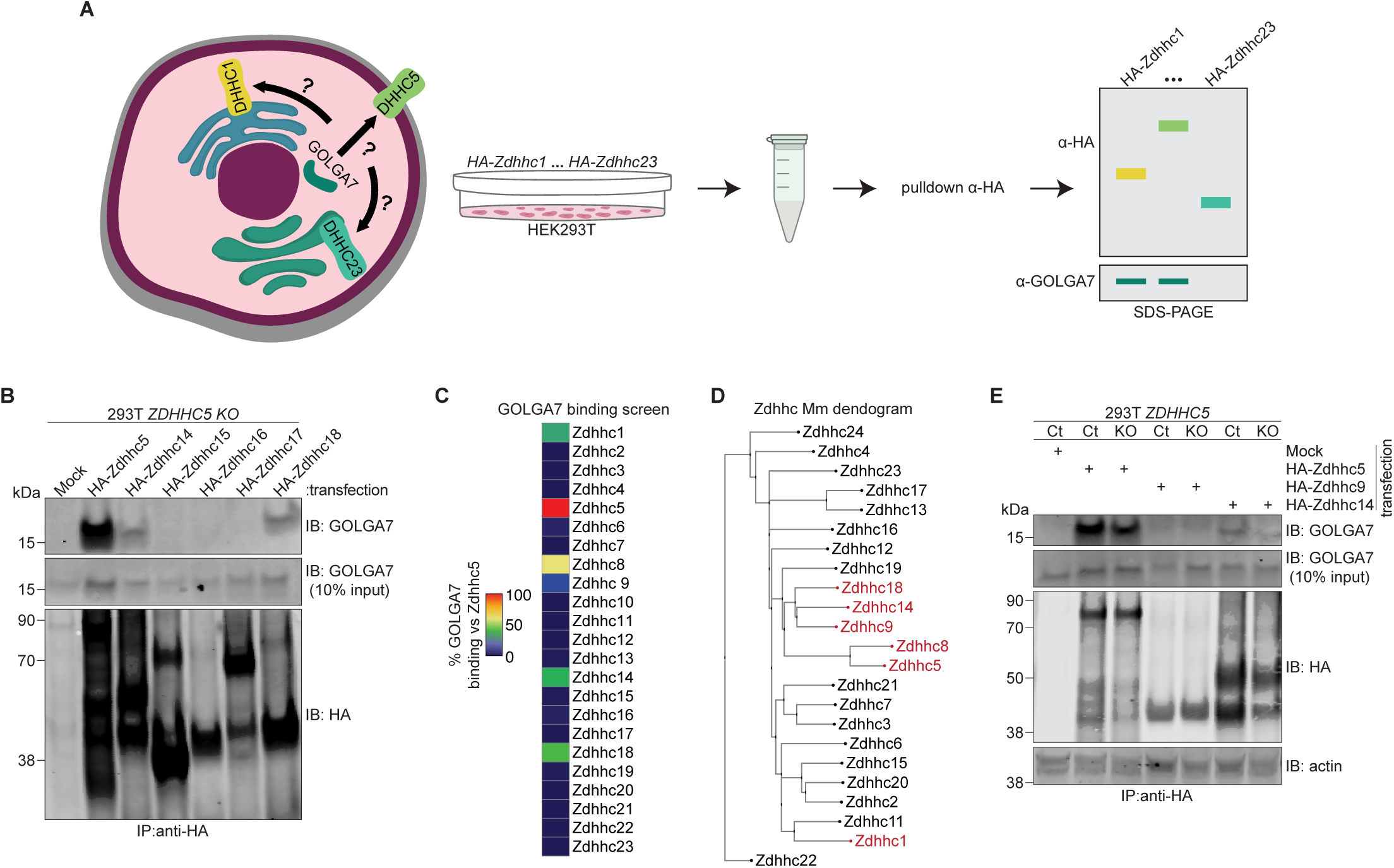
GOLGA7 interacts with six mammalian ZDHHC enzymes. (A) Schematic illustrating co-immunoprecipitation screen of all mammalian ZDHHC enzymes for endogenous GOLGA7 binding. (B) Representative co-immunoprecipitation analysis of endogenous GOLGA7 binding in a *ZDHHC5* gene-disrupted (”KO”) cell line transfected as indicated. (C) Heatmap representing percent GOLGA7 binding to each tested mammalian Zdhhc-family enzyme, based on western blot quantification, normalized to Zdhhc5 binding. (D) Phylo.io-generated neighbor-joining tree of all mouse Zdhhc enzymes based on sequence conservation. (E) Co-immunoprecipitation analysis of endogenous GOLGA7 binding in Control (Ct) and *ZDHHC5* gene disrupted (”KO”) cell lines transfected as indicated. Blot is representative of three independent experiments.

In *ZDHHC5^KO^* cells, endogenous GOLGA7 was recovered in co-IPs from cells expressing Zdhhc1, Zdhhc5, Zdhhc8, Zdhhc9, Zdhhc14 and Zdhhc18, with the highest recovery observed with Zdhhc5. We compared this interaction pattern to the overall sequence similarity between all 23 Zdhhc proteins using a neighbor-joining tree (**Fig. 1*D***). Zdhhc5, Zdhhc8, Zdhhc9, Zdhhc14, and Zdhhc18 were all found within a common branch, correlating with their ability to complex with GOLGA7. Unexpectedly, Zdhhc19 also clustered within this branch but was the only PAT in the cluster that failed to complex with GOLGA7 (**Fig. 1*C*, Supplementary Fig. S1*B***). By contrast, Zdhhc1 did complex with endogenous GOLGA7 despite sharing little sequence conservation with Zdhhc5 outside the cysteine rich domain (CRD), DHHC active site, and two short motifs immediately downstream of the active site (discussed further below). Zdhhc1 therefore occupied a distant site on the tree from the Zdhhc5 cluster (**Fig. 1*D***). Similar patterns of complex formation were observed in 293T Control cells, with Zdhhc14 showing higher recovery of endogenous GOLGA7 compared to *ZDHHC5^KO^* cells, and a loss of detectable GOLGA7 recovery by Zdhhc1 (**Fig. 1*E*, Supplementary Fig. S1*C***). In sum, these data suggest that GOLGA7 is more promiscuous than previously thought, capable of forming complexes with at least six distinct ZDHHC proteins.

### An RNYR motif is required for ZDHHC interactions with GOLGA7

The six mouse Zdhhc proteins that formed complexes with GOLGA7 were generally as similar at the amino acid sequence level to their human orthologs (94 ± 4%) as the Zdhhc proteins that failed to complex with GOLGA7 (89 ± 11%) (Mann-Whitney U, p > 0.05, **Supplementary Fig. S1*A***). Thus, global differences in amino acid sequences divergence did not appear to account for differences in complex formation. To investigate whether more specific sequence features accounted for Zdhhc-GOLGA7 complex formation, we pursued a two-pronged approach combining deletion mapping and computational inference.

First, we generated a series of Zdhhc5 deletion mutants lacking various regions of the protein, including the N-terminus and transmembrane (TM) domains (HA-Zdhhc5^ΔN-term^), the cytosolic loop containing the enzyme DHHC active site and cysteine rich domain (Zdhhc5^ΔDHHC^), or the C-terminus (HA-Zdhhc5^ΔC-term^) (**Fig. 2*A***). Note that Zdhhc5 and other DHHC enzymes with long C-termini are prone to proteolytic degradation, which results in multiple bands on immunoblot. Additionally, Zdhhc5 and other GOLGA7 binders exhibit decreased stability and a propensity for oligomerization or aggregation when not occupied by GOLGA7 (20), sometimes resulting in an SDS-resistant crosslinked ladder on immunoblot (**Supplementary Fig. 2D**). Co-IP experiments demonstrated that endogenous GOLGA7 was recovered from 293T cells overexpressing full-length Zdhhc5 and, to a lesser degree, HA-Zdhhc5^ΔC-term^ (**Fig. 2*B*)**. Consistent with these biochemical results, expression of full-length HA-Zdhhc5, and to a lesser degree HA-Zdhhc5^ΔC-term^, restored CIL56-induced cell death in 293T^N^ (stably expressing nuclear-localized mKate2) *ZDHHC5^KO^* cells. However, overexpression of HA-Zdhhc5^ΔN-term^ and HA-Zdhhc5^ΔDHHC^ failed to restore CIL56 lethality (**Fig. 2C**). Despite the lower expression of HA-Zdhhc5^ΔDHHC^ protein expression (**Fig. 2B**), these results indicate that the cytosolic loop of Zdhhc5, which was absent from both non-binding mutants, is essential for the interaction of Zdhhc5 with endogenous GOLGA7 in 293T cells.

**Figure 2.**
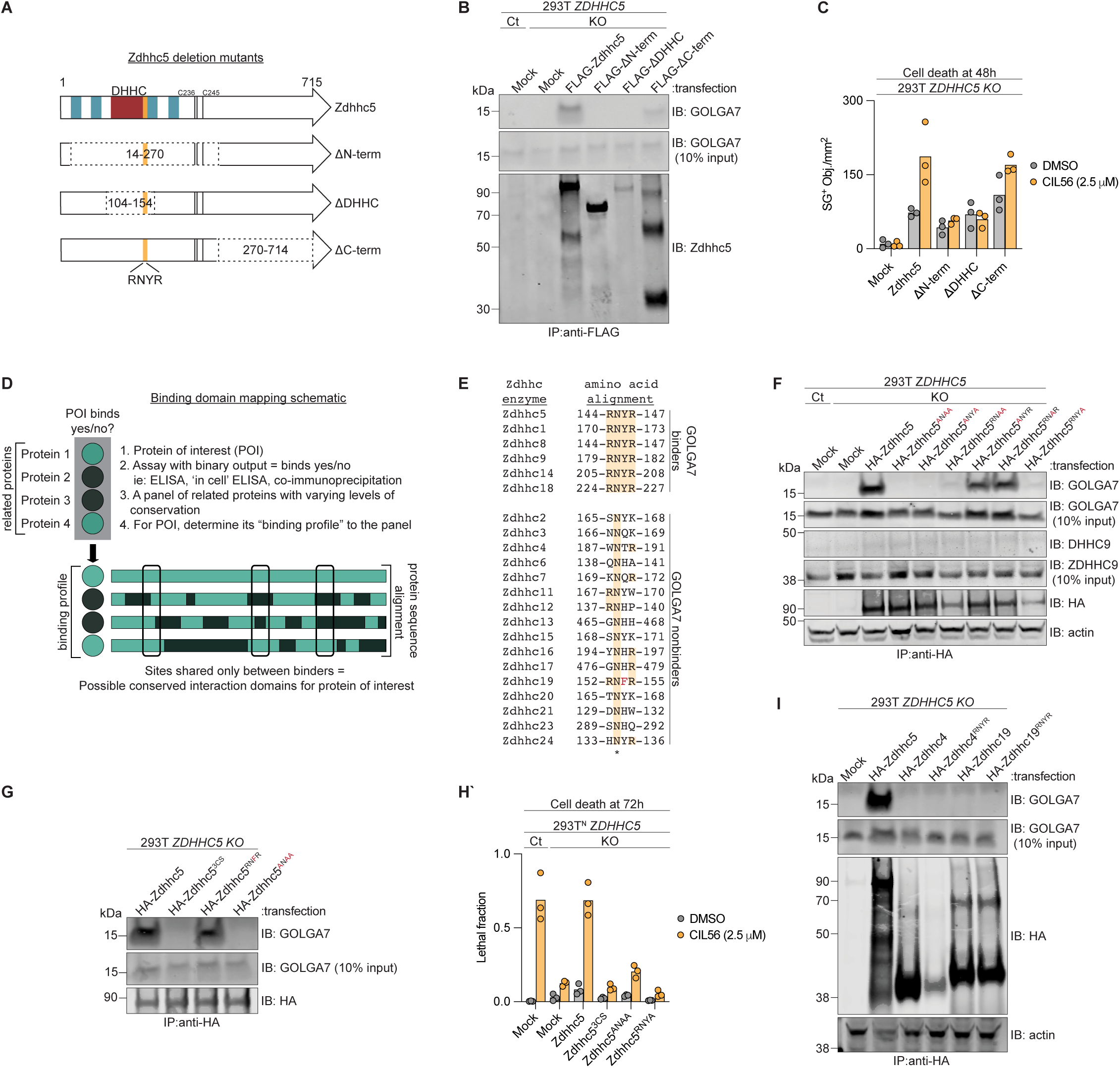
RNYR motif is a key interaction domain on GOLGA7 binders. (A) Schematic of Zdhhc5 deletion mutants. Blue bars represent transmembrane domains, black bars indicate acylated cysteines, and red represents the DHHC active site and conserved cysteine rich domain. (B) Co-immunoprecipitation analysis of endogenous GOLGA7 binding in Control (Ct) and *ZDHHC5* gene disrupted (”KO”) cell lines transfected as indicated. (C) Cell death analysis of *ZDHHC5^KO^* cell lines transfected as indicated. (D) Schematic of motif mapping strategy for interaction domain identification. Green indicates the protein of interest (POI) binder in this schematic. (E) Local amino acid alignment of all mouse Zdhhc-family proteins, highlighting the RNYR motif uniquely shared by GOLGA7 binders. (F-G) Co-immunoprecipitation analysis of GOLGA7 binding in Ct and *ZDHHC5^KO^*cell lines transfected as indicated. (H) Cell viability analysis of Ct and *ZDHHC5^KO^* cell lines transfected with constructs corresponding to (F) and (G). (I) Co-immunoprecipitation analysis of endogenous GOLGA7 and ZDHHC9 binding in Ct and *ZDHHC5^KO^* cell lines transfected as indicated. (B) and (F) are representative of two independent experiments. (G) and (H) are representative of four independent experiments. Data are in (C) and (I) are from three independent experiments; each datapoint represents one independent replicate.

Protein-protein interactions often rely on short linear motifs within one or both binding partners (22–24). We hypothesized that specific amino acid motifs may be essential for the binding of Zdhhc proteins to GOLGA7. Accordingly, we searched for conserved amino acid motifs greater than one residue in length that were found in Zdhhc-GOLGA7 interactors (Zdhhc1, Zdhhc5, Zdhhc8, Zdhhc9, Zdhhc14 and Zdhhc18) but absent from the non-interacting set (**Fig. 2*D***). This analysis identified a single motif that was unique to GOLGA7 binders: RNYR (^144^RNYR^147^ in Zdhhc5). This motif lies within the cytosolic loop, ten residues C-terminal to the DHHC motif, and the region defined by deletion mapping as essential for interaction with endogenous GOLGA7 (**Fig. 2*E*, Supplementary Fig. S2*A***). Thus, we hypothesized that the RNYR motif was essential for Zdhhc-GOLGA7 complex formation.

To experimentally test whether the RNYR motif is required for Zdhhc complex formation with GOLGA7, we generated a series of alanine substitution mutants in Zdhhc5, the strongest interactor. Single alanine substitutions of Arg144 or Tyr146 (“ANYR” or “RNAR”, respectively) did not disrupt interaction, as both mutants retained the ability to co-IP endogenous GOLGA7 and expressed at similar levels to wild-type protein (**Fig. 2*F*)**. However, “ANYA” and “RNAA” double mutants failed to complex with GOLGA7. Notably, the fourth residue in the RNYR motif, Arg^147^, appeared to be essential for GOLGA7 binding, as a single Arg147Ala mutation (“RNYA”) abolished interaction with endogenous GOLGA7 despite effective protein expression (**Fig. 2*F*).** Consistent with these results, a triple alanine mutant (Zdhhc5^ANAA^), which retained only the universally conserved asparagine residue, failed to co-IP endogenous GOLGA7 (**Fig. 2*G***). Mirroring the co-IP results, sensitivity to CIL56-induced cell death was restored in 293T ZDHHC5^KO^ cells by overexpression of wild-type HA-Zdhhc5 but not the triple alanine mutant HA-Zdhhc5^ANAA^ or the single point mutant HA-Zdhhc5^RNYA^ (**Fig. 2*H***).

The Zdhhc5^RNYR^ motif mutants could fail to complex with GOLGA7 and thus fail to facilitate CIL56 cell death due to either a loss of direct interaction or because this mutant cannot traffic to the region of the cell necessary for complex formation with Zdhhc5. For example, a previously identified Zdhhc5 mutant that cannot be *S*-acylated on three cysteine residues (Zdhhc5^3CS^), cannot reach the plasma membrane and accumulates on internal membranes (2). Using 293T *ZDHHC5^KO^* cells, we confirmed the aberrant accumulation of the Zdhhc5^3CS^ mutant throughout the cytoplasm surrounding the nucleus; by contrast, wild-type Zdhhc5 and the most disruptive mutant RNYR motif mutant (Zdhhc5^ANAA^) both localized predominantly to the plasma membrane, as determined by colocalization with the plasma membrane marker Cadherin (**Supplemental Fig S2*D-H***). Thus, the RNYR motif does not appear to drive Zdhhc5 intracellular localization but instead presumably affected complex formation and cell death through a more proximate disruption of protein interactions.

Our results thus far demonstrated that the Zdhhc RNYR motif is necessary for complex formation with GOLGA7. To determine whether this motif was sufficient for GOLGA7 complex formation, we installed a RNYR motif at the corresponding site in GOLGA7 non-binding proteins, namely Zdhhc4 (^189^WNTR^192^ ➔ ^189^RNYR^192^). We also mutated the closely related motif found in the GOLGA7 non-binder Zdhhc19 (^152^RNFR^155^) to the more optimal motif found in all GOLGA7 binders, ^152^RNYR^155^. In both cases, installation of the RNYR motif did not enable co-IP of endogenous GOLGA7 when these proteins were expressed in 293T *Zdhhc5^KO^* cells (**Fig. 2*I***). Of note, mutation of RNYR to RNFR in Zdhhc5 (to mimic the sequence of Zdhhc19 at this motif) did not eliminate endogenous GOLGA7 interaction, as assessed by co-IP (**Fig. 2*G***). As a negative control, we utilized the Zdhhc5 acylation-deficient mutant (Zdhhc5^3CS^), previously shown to be incapable of complex formation with GOLGA7 (2) (**Fig. 2*G***). Thus, the Zdhhc RNYR motif appeared necessary but not sufficient for directing Zdhhc complex formation with GOLGA7.

### A GOLGA7 motif required for ZDHHC5 complex formation

Complex formation between ZDHHC5 and GOLGA7 is essential for cell death induced by CIL56 in 293T, HT-1080, and other cancer cell lines (2). However, ZDHHC5 is also reported to form complexes with GOLGA7B, a paralogous protein that shares 79% primary sequence identity with GOLGA7 (17) (**Fig. 3*A***). Whether GOLGA7 and GOLGA7B serve interchangeable functions in the regulation of CIL56-induced cell death and, by extension, ZDHHC5 function, was unclear. To investigate this, we performed co-IP assays in our established 293T *GOLGA7^KO^* cells (2) transfected with FLAG-tagged GOLGA7B or GOLGA7. Overexpressed 3xFLAG-GOLGA7B pulled down endogenous ZDHHC5, as did the positive control constructs 1xFLAG-GOLGA7 and 3xFLAG-GOLGA7, the latter yielding a stronger anti-FLAG signal by immunoblot. Given its higher expression, 3xFLAG-GOLGA7 and was used in subsequent experiments (**Fig. 3*B***).

**Figure 3.**
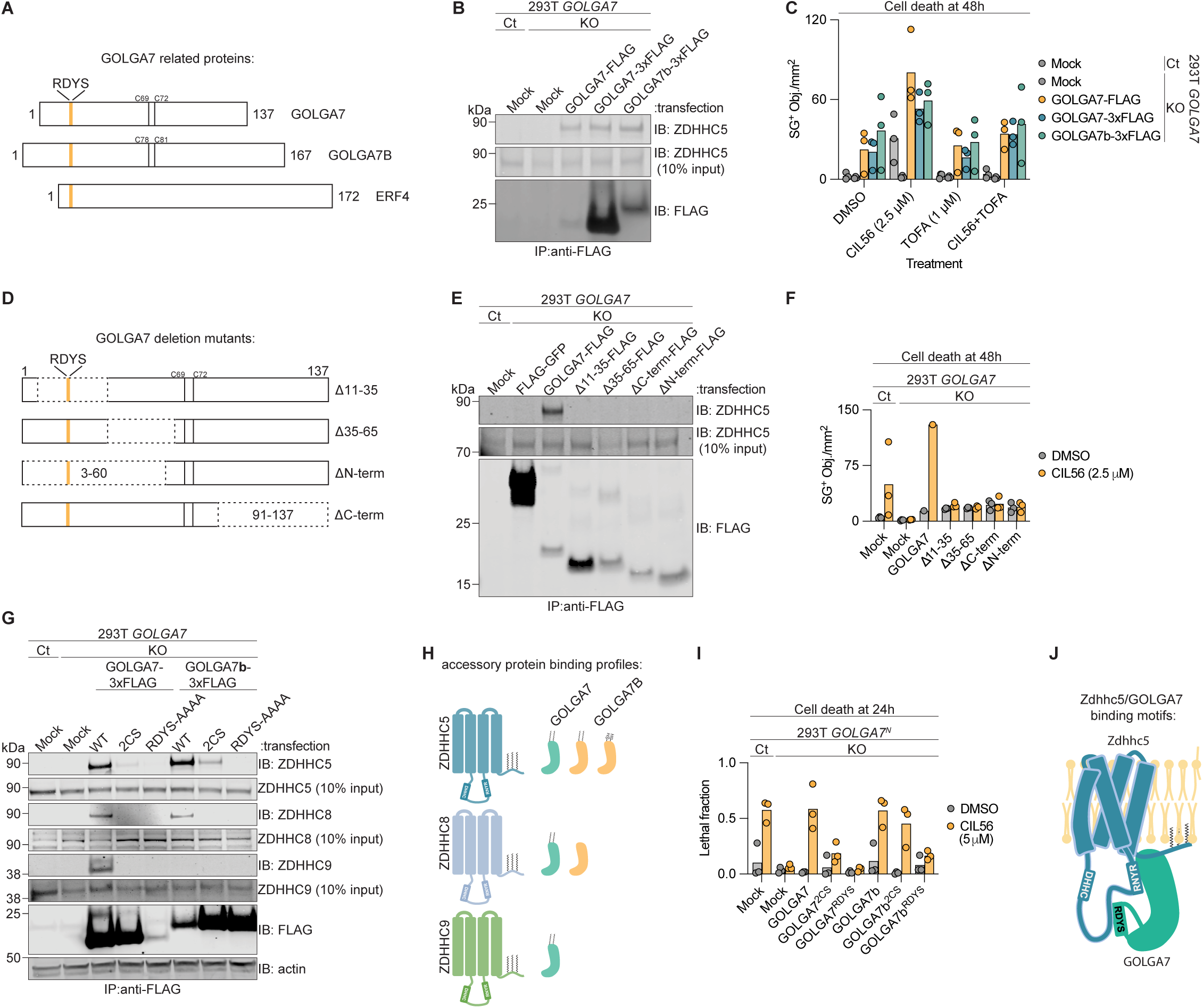
RDYS is a key ZDHHC interaction domain shared by GOLGA7, GOLGA7b, and Erf4. (A) Schematic of GOLGA7, GOLGA7B, and Erf4 structures indicating the conserved RDYS motif (orange) and acylated cysteines (black). (B) Co-immunoprecipitation analysis of endogenous ZDHHC5 binding from Control (Ct) and *GOLGA7* gene-disrupted (”KO”) cell lines transfected as indicated. Note: the 3xFLAG tag improves GOLGA7 detection. Blot is representative of two independent experiments. (C) Cell death analysis of cells transfected as in (B) and treated with CIL56 (2.5 μM) or TOFA (1 μM) as indicated. (D) Schematic of GOLGA7 deletion mutants with acylated cysteines in black. (E) Co-immunoprecipitation analysis of endogenous ZDHHC5 binding in Ct and *GOLGA7^KO^* cells transfected as indicated. Blot is representative of two independent experiments. (F) Cell death analysis of cells transfected as in (E). (G) Co-immunoprecipitation analysis of endogenous ZDHHC5, ZDHHC8, and ZDHHC9 binding in Ct and *GOLGA7^KO^* cells transfected as indicated. Blot is representative of four independent experiments. (H) Schematic summarizing data from (G), with GOLGA7 in seafoam green and GOLGA7B in orange. (I) Cell viability analysis of Ct and *GOLGA7^KO^*NLR cells transfected as in (G). (J) Schematic of the Zdhhc5 (teal) and GOLGA7 (green) complex at the plasma membrane, showing respective binding motifs and the DHHC active site. (C), (F), and (I) data are each from three independent experiments, except for the wild-type positive control in (F), which is from one replicate.

The ability of GOLGA7 and GOLGA7B to promote CIL56-induced cell death correlated with their complex formation with ZDHHC5. Overexpression of either GOLGA7 or GOLGA7B in 293T *GOLGA7^KO^* cells restored cell death in response to CIL56 (**Fig. 3*C***). An inhibitor of CIL56-induced cell death, TOFA (2), suppressed cell death across all conditions, indicating that GOLGA7 or GOLGA7B overexpression did not change the mode of cell death in response to CIL56. Thus, GOLGA7 and GOLGA7B can in principle function interchangeably to promote CIL56-induced cell death. Note that normally most cells appear to express GOLGA7 and not GOLGA7B, explaining why disruption of GOLGA7 alone is able to fully inhibit cell death in response to CIL56 (17). Nevertheless, we reasoned that we could use the shared ability of GOLGA7 and GOLGA7B to reconstitute cell death in 293T *GOLGA7^KO^* cells for subsequent analyses to determine key residues necessary for complex formation with Zdhhc5.

To identify determinants of ZDHHC-GOLGA7 complex formation, we designed and tested a series of GOLGA7 deletion mutants encompassing distinct regions of the protein: Gly^10^-Glu^35^ (“Δ11-35”), Glu^35^-Tyr^65^ (“Δ35-65”), Pro^3^-Leu^60^ (“ΔN-term”), and Lys^91^-Arg^137^ (“ΔC-term”) (**Fig. 3*D***). We overexpressed these proteins in 293T Control and *GOLGA7^KO^* cells and performed anti-FLAG immunoprecipitation, using 3xFLAG-GOLGA7 as a positive control. While 3xFLAG-GOLGA7 co-immunoprecipitated endogenous ZDHHC5 as expected, none of the four GOLGA7 mutants recovered detectable levels of endogenous ZDHHC5, despite reasonably equal expression of all proteins (**Figure 3E**). These biochemical data mirrored functional data, which indicated that none of the GOLGA7 mutants restored CIL56-induced cell death when expressed in 293T^N^ *GOLGA7^KO^*cells (**Fig. 3*F***).

Given that both GOLGA7 and GOLGA7B complex with ZDHHC5 and promote CIL56-induced cell death, we hypothesized that they shared one or more conserved motifs necessary (but not sufficient) for interaction with ZDHHC proteins. Multiple sequence alignment between GOLGA7, GOLGA7B and the *S. pombe* accessory protein Erf4, identified a conserved four amino acid motif, ^16^RDYS^19^ in GOLGA7 and ^25^RDYS^28^ in GOLGA7 (**Fig. 3*A***). Unlike the corresponding wild-type proteins, a GOLGA7 quadruple-alanine mutant (RDYS ➔ AAAA, “GOLGA7^AAAA^”) failed to co-IP endogenous ZDHHC5 in 293T *GOLGA7^KO^* cells, despite effective protein expression (**Fig. 3*G***). This resembled the phenotype of an *S*-acylation-defective Cys-to-Ser mutant (GOLGA7^2CS^) we identified previously (2) and used as a negative control (**Fig. 3G**). A corresponding GOLGA7B^AAAA^ quadruple alanine mutant similarly failed to recover endogenous ZDHHC5 (**Supplementary Fig. S3*A***). Unlike wild-type GOLGA7 and GOLGA7B, the GOLGA7^AAAA^ and GOLGA7B^AAAA^ mutants also failed to restore sensitivity to CIL56-induced cell death when overexpressed in 293T^N^ *GOLGA7^KO^* cells, much like the acylation-deficient GOLGA7^2CS^ control (2) (**Fig. 3*I***). Thus, the RDYS motif, conserved between GOLGA7 and GOLGA7B, is essential for functional interaction with ZDHHC5. Unlike the Cys residues disrupted in the GOLGA7^2CS^ mutant, the RYDS motif was not obviously related to protein *S*-acylation, leading us to hypothesize that it could be important in some way for complex formation between GOLGA7/B and Zdhhc proteins.

### Unique functional determinants of GOLGA7B interaction with ZDHHC5

Despite the shared RDYS-dependent binding of GOLGA7 and GOLGA7B to ZDHHC5, their interactions were not identical. We previously determined that GOLGA7 palmitoylation is likely required for interaction with ZDHHC5, as demonstrated using the *S*-acylation-deficient GOLGA7^2CS^ mutant (2). To investigate whether GOLGA7B also depends on *S*-acylation for ZDHHC5 interaction, we generated an equivalent *S*-acylation-deficient 3xFLAG-GOLGA7B mutant (GOLGA7B^2CS^). Co-IP experiments demonstrated that GOLGA7B^2CS^ expressed in *GOLGA7^KO^*cells effectively recovered some endogenous ZDHHC5, albeit lower amounts than wild-type GOLGA7B (**Fig. 3, *G*** and ***H***). Functionally, overexpression of GOLGA7B^2CS^ also restored CIL56 lethality in 293T^N^ *GOLGA7^KO^* cells to similar levels as wild-type GOLGA7B (**Fig. 3*I***). These results suggest that, unlike GOLGA7, *S*-acylation at two orthologous cysteine residues in GOLGA7B may not be necessary for interaction with ZDHHC5 or for the induction of non-apoptotic cell death.

Given that GOLGA7 could complex with approximately one-quarter of all mammalian ZDHHC enzymes (**Fig. 1*C***), and that both GOLGA7 and GOLGA7B interact with ZDHHC5, we hypothesized that GOLGA7B might also interact with other ZDHHC-family proteins in an RYDS-dependent manner. Indeed, co-IP experiments demonstrated that 3xFLAG-GOLGA7B pulled down endogenous ZDHHC8 in addition to ZDHHC5 (**Fig. 3, *G*** and ***H***). However, we also noted important differences in the behavior of GOLGA7 and GOLGA7B in relation to complex formation with Zdhhc-family proteins. First, wild-type GOLGA7 but not wild-type GOLGA7B could complex with ZDHHC9 in co-IP studies (**Fig. 3*G***). Second, the *S*-acylation deficient (2CS) mutant of GOLGA7B could complex to some degree with ZDHHC5 but not at all with ZDHHC8 (**Fig. 3, *G*** and ***H***). Thus, GOLGA7 and GOLGA7B are not identical in their ability to form complexes with specific ZDHHC proteins (**Fig. 3*H***). Nevertheless, our mapping approaches identified conserved motifs on both Zdhhc enzymes and on GOLGA7/B accessory proteins that appeared necessary for complex formation and the induction of non-apoptotic cell death (**Fig. 3*J***).

### Structural analysis of the Zdhhc5-GOLGA7 interaction

We hypothesized that residues identified above through biochemical and phenotypic analysis as necessary for Zdhhc5-GOLGA7 complex formation were at critical interfaces necessary for physical interaction between these two proteins. To test this hypothesis, we used cryogenic electron microscopy (cryo-EM) to experimentally determine the structure of the Zdhhc5-GOLGA7 complex. We obtained a 3D reconstruction at an estimated local resolution ranging from 3.9–6 Å. We then used AlphaFold 3 (25) to predict a structure for this complex, referred to here as the “reference” structure (Zdhhc5-GOLGA7_AF3_, see more below). We then combined the experimentally determined and reference structures to construct a hybrid structural model of the Zdhhc5-GOLGA7 complex (**Fig. 4*A***). For comparison, we also generated an AlphaFold 3 (25) model of human ZDHHC5 in complex with GOLGA7 and found that this model closely matched the reference structure (**Supplementary Fig. S*4***). In the hybrid structure, the backbone of Zdhhc5 and GOLGA7 were modeled using the cryo-EM density, and side chains were manually modeled where resolvable. For residues where side chain density was not interpretable, we retained side chain conformations predicted by AlphaFold 3. This hybrid cryo-EM model allowed us to visualize the full backbone architecture and key side chain interactions at the Zdhhc5-GOLGA7 interface, while incorporating predicted elements where this was justified by lack of density.

**Figure 4.**
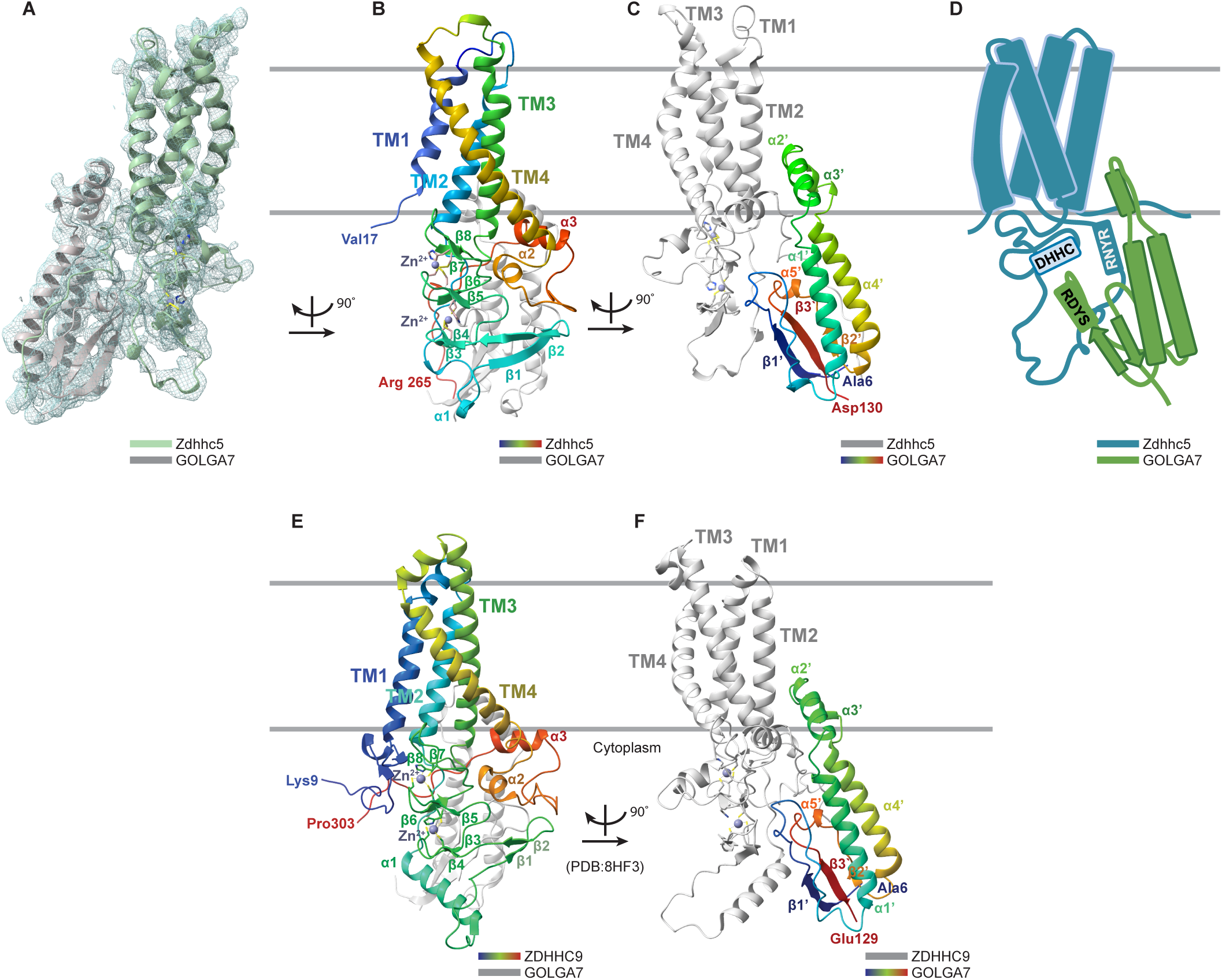
Overview of Zdhhc5/GOLGA7 and comparison with ZDHHC9/GOLGA7. (A) Cryo-EM density map of the Zdhhc5-GOLGA7 complex overlaid with the hybrid model (Zdhhc5 in light green, GOLGA7 in grey). (B) Ribbon diagram of Zdhhc5– GOLGA7 model with Zdhhc5 colored from blue (N-terminus) to red (C-terminus) for each chain, and GOLGA7 in grey. Grey lines represent estimated plasma membrane boundaries. (C) Ribbon diagram of Zdhhc5–GOLGA7 model with GOLGA7 colored from blue (N-terminus) to red (C-terminus), and Zdhhc5 in grey. (D) Schematic summary of the Zdhhc5/GOLGA7 complex, indicating relative positions of the DHHC active site, RNYR and RDYS motifs, and membrane orientation. (E) and (F) Ribbon diagram of ZDHHC9/GOLGA7 complex (PDB 8HF3, from (18)) shown with the same orientation and coloring as in (B) and (C) respectively, for comparison.

In the Zdhhc5-GOLGA7_AF3_ reference model, the structured regions of Zdhhc5 were predicted with very high accuracy. AlphaFold 3 predicted local distance difference test (pLDTT) scores, a per-atom confidence estimate on a 0-100 scale (where higher values indicate higher confidence), were high (> 90) for all structured elements of the complex. Flexible loops and helical termini exhibited moderate confidence (90 > pLDTT > 70), apart from the unstructured N-terminal residues and the least structured loop region between TM2 and TM3 (residues ∼76-90), which ranged from confident to very low (< 50). The C-terminus of ZDHHC5 (residues 260-715) was not modeled. The predicted template modeling (pTM) and the interface predicted template modeling (ipTM) scores are additional metrics of the accuracy of protein structure. The pTM score for Zdhhc5-GOLGA7_AF3_ was 0.5, where 0.5 or above indicates the overall predicted fold for the complex is a reasonable prediction of the native structure (25–27). The ipTM measures the accuracy of the predicted relative position between subunits in a complex. The ipTM for Zdhhc5-GOLGA7_AF3_ was 0.89, where values > 0.8 represent confident predictions (25–27). While the AlphaFold 3 model provided a reliable starting point for modeling from our cryo-EM map, manual adjustment of backbone geometry was required during model building, particularly for the orientation of transmembrane helices, to better fit the density (**Fig *4A**, B, C***). To summarize the architecture of the Zdhhc5– GOLGA7 complex, we generated a simplified cartoon highlighting the membrane topology, key helices, and the spatial positioning of the RNYR and RDYS motifs within the assembled structure (**Fig. 4*D***).

Overlaying our cryo-EM-based Zdhhc5-GOLGA7 model with a recently reported cryo-EM structure for the ZDHHC9-GOLGA7 complex (PDB: 8HF3) (20) revealed high structural similarity (**Fig. *4E, F***). A TM score was used to assess the topological similarities between Zdhhc5-GOLGA7 and ZDHHC9-GOLGA7. Structural alignment using Foldseek online (28) predicted a tTM score (TM score normalized to the target length) of 0.82, where scores of 0.5 to 1.0 indicate similar fold or family (26). Both complexes contain four transmembrane (TM) helices, a large cytosolic loop between TM2 and TM3, and a cytosolic domain following TM4 that precedes the long C-terminal tail (**Fig. 4*B, E*)**. The cytosolic loop features one α-helix and four anti-parallel β-sheets that together form two zinc finger motifs stabilizing the DHHC active site. Additionally, two conserved α-helices are present in the cytosolic region following TM4. In both models, GOLGA7 has two α helices (α2’ and α3’) inserted into the membrane in a position similar to the α5 helix unique to the ZDHHC family member ZDHHC20 (18) (**Fig. 4*C, F***). This helix in ZDHHC20 plays a significant role in its autoacylation activity, reducing the rate by about half when mutated (9). The absence of this α-5 equivalent helix in Zdhhc5 and ZDHHC9 may underlie their dependency on accessory proteins such as GOLGA7 for full function.

### Comparisons with the ZDHHC9-GOLGA7 complex structure

Our model and the previously determined ZDHHC9 model (PDB: 8HF3) (20) indicated that both DHHC enzymes engage GOLGA7 through four primary interfaces. In our Zdhhc5-GOLGA7 hybrid model, the Zdhhc5 RNYR and GOLGA7 RDYS motifs were positioned within two distinct interfaces. Mutations in the Zdhhc5 RNYR motif (RNYR to ANAA or RNYA) disrupted binding, with the second arginine (Arg^147^) being essential for co-IP interaction (**Fig. 2*G***). In our structural model (**Fig. 5*A***), Zdhhc5 Arg^147^, located on TM3, interacted with the α2’ and α3’ helices of GOLGA7 (**Fig. 5*B***). Side chains were consistent with a cation-π interaction between Zdhhc5 Arg^147^ and the aromatic ring of Tyr^76^ (**Fig. 5*B***). These structural insights were fully concordant with the importance of Arg^147^ for complex formation discovered through biochemical and functional assays.

**Figure 5.**
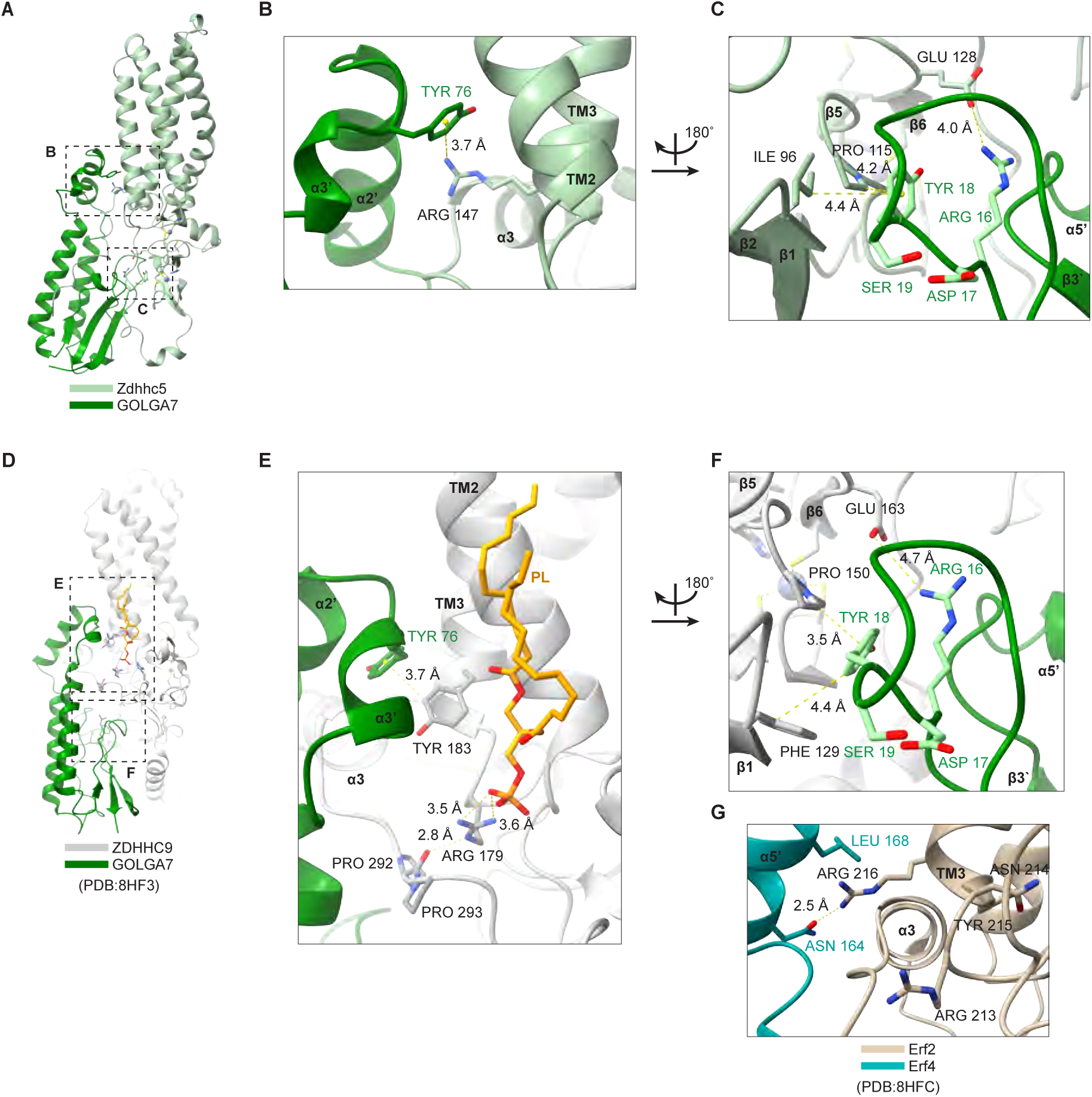
Key residues at the ZDHHC-GOLGA7 interaction surface. (A) The Zdhhc5-GOLGA7 complex (Zdhhc5 in light green, GOLGA7 in dark green). (B) Arg^147^ from the RNYR motif (TM3) interacts with the α2’ helix of GOLGA7 at Tyr^76^. Hydrogen bonds are represented by dashed, yellow lines. (C) GOLGA7 RDYS motif residues Arg^16^ and Tyr^18^, located in the loop following the β1’ strand, interact with Ile^96^, Pro^115^ and Glu^128^ the zinc finger region. (D) The ZDHHC9-GOLGA7 complex (ZDHHC9 in grey, GOLGA7 in dark green, PDB 8HF3, from (18)). (E) Arg^179^ (RNYR motif) helps coordinate a phospholipid (yellow) and the PPII helix of ZDHHC9, which in turn docks into GOLGA7. Tyr^183^ (TM3) stacks with Tyr^76^ on the GOLGA7 α3’ helix. (F) GOLGA7 RDYS residues Arg^16^ and Tyr^18^, in the loop following the β1’ strand, interact ZDHHC9 Phe^129^, and Pro^150^ and Glu^165^ within zinc finger domains. (G) In the *S. cerevisiae* Erf2-Erf4 complex (PDB: 8HFC (18)), R^216^ (RNYR motif, TM3) forms a hydrogen bond with Erf4 Asn^164^ and interacts with Leu^168^ on the α5’ helix. Blue indicates nitrogen and red indicates oxygen across all panels.

The GOLGA7 ^16^RDYS^19^ motif was likewise critical for complex formation with ZDHHC-family proteins. In our structural model, the GOLGA7 RDYS motif was located on a short loop following the β1′ strand and interfaces with the Zdhhc5 zinc finger motifs that stabilized the active DHHC site. While the residues on this loop were not sufficiently resolved in our cryo-EM map, structural comparison to our reference model suggested that the first residue of the GOLGA7 RDYS motif (Arg^16^) likely interacted with Zdhhc5 Glu^128^ through a charge-charge interaction, potentially forming a salt bridge (**Fig. 5*C***). Additionally, GOLGA7 Tyr^18^ was positioned to interact with Zdhhc5 Pro^115^ through CH-π stacking interactions and may also form weaker interactions, such as transient van der Waals contacts, with Zdhhc5 Ile^96^. Together, these interactions suggested that GOLGA7 Arg^16^ and Tyr^18^ within the RDYS motif helped stabilize the active site conformation through binding to the zinc finger domain, consistent with their functional importance.

The structural importance of the Zdhhc5 RNYR and the GOLGA7 RDYS motifs was conserved in the ZDHHC9-GOLGA7 complex (PDB: 8HF3) (20) (**Fig. 5*D***). ZDHHC9 Arg^179^ (<u>R</u>NYR) helped form a positively charged patch that coordinated a lipid-like density, proposed to phosphatidic acid (18) (**Fig. 5*E***). Given the structural conservation between ZDHHC9-GOLGA7 and Zdhhc5-GOLGA7, Zdhhc5 may similarly bind this lipid, though lipid density was unresolvable in our map. In ZDHHC9-GOLGA7, this interaction stabilized the TM2 and TM3 and type II polyproline (PPII) helices of ZDHHC9 and its interaction with GOLGA7. Tyr^183^ (RNYR<u>Y</u>) on ZDHHC9 TM3 directly interacted with GOLGA7 Tyr^76^ on α3’ helix thru π-π stacking (**Fig. 5*E***), an interaction shown to be functionally important as mutation of Tyr^76^ decreased ZDHHC9 catalytic activity (18). As in the Zdhhc5-GOLGA7 structure, the GOLGA7 RDYS site also appeared to lie at a key interaction surface in the ZDHHC9-GOLGA7 structure at the ZDHHC9 zinc finger domains. GOLGA7 Arg^16^ interacted with ZDHHC9 Glu^163^ through charge-charge interaction, and Tyr^18^ forms π-π and CH-π stacking interactions with ZDHHC9 Phe^129^ and Pro^150^, respectively (**Fig. 5*F***).

Despite the evolutionary divergence between yeast and human DHHC enzymes (only 31% identity between Erf2 and ZDHHC9) (**Supplementary Fig. S*5*)**, the key residue of the RNYR motif, Arg216 on Erf2 TM3, also formed an interactive domain with Erf4, the yeast GOLGA7 ortholog (**Fig 5*G***). The guanidinium group of Arg^216^ donated a hydrogen bond to the side-chain carbonyl oxygen of Erf4 Asn^164^ on the α5’ helix and may further stabilize the interaction via hydrophobic contacts with the aliphatic side chain of Erf4 Leu^168^. Thus, the structural importance of the RNYR motif appears conserved across species, from yeast to humans.

## DISCUSSION

CIL56 is a lethal small molecule that induces an unusual form of non-apoptotic cell death dependent on the ZDHHC5-GOLGA7 protein complex (2, 3). How this complex promotes cell death in response to CIL56 remains to be clarified. One possibility is that CIL56 directly activates the ZDHHC5-GOLGA7 complex, such that deletion of either *ZDHHC5* or *GOLGA7*, or disruptions to protein complex formation, are sufficient to inhibit cell death. Alternatively, the ZDHHC5-GOLGA7 complex may *S*-acylate one or more proteins in a manner that becomes lethal upon CIL56 treatment, for instance by impairing protein trafficking (2). Regardless, here we could use CIL56 as a probe, leveraging resistance to cell death as a functional readout to help identify key residues necessary for the ZDHHC5-GOLGA7 complex itself. Remarkably, amino acid residues identified through this approach, including the Zdhhc5 RDYS motif, were subsequently validated as essential for complex formation with GOLGA7 in our CryoEM and AlphaFold 3 structures. This strategy may prove generally useful for mapping important amino acids in proteins involved in this and other forms of non-apoptotic cell death.

We find that assembly of the ZDHHC5-GOLGA7 complex is mediated by specific short motifs and individual amino acid residues on both proteins. Within the ZDHHC5 RNYR motif, mutation of R^147^ to Ala (“RNYA”) alone was sufficient to completely disrupt co-immunoprecipitation with endogenous GOLGA7, despite robust expression of this mutant. The finding is rationalized by our Zdhhc5-GOLGA7 model, where Zdhhc5 R^147^ appears to contribute directly to the interaction interface. In addition to Zdhhc5, GOLGA7 also interacted to some degree with Zdhhc1, Zdhhc8, Zdhhc9, Zdhhc14 and Zdhhc18. We expect that these interactions to require the RNYR motifs, but this remains to be experimentally verified. Zdhhc1, Zdhhc8, Zdhhc9, Zdhhc14, and Zdhhc18 localize to distinct subcellular compartments and function in growth factor signaling, immune signaling, and stem cell function (10–12, 29, 30). Thus, GOLGA7 may play a broad role in regulating protein *S*-acylation within the cell, and disruption of GOLGA7 may impair the function of multiple ZDHHC enzymes. At the same time, the ability of GOLGA7 and its paralog GOLGA7B to functionally substitute for one another in some ZDHHC complexes raises the possibility of compensatory regulatory mechanisms.

While GOLGA7 exhibits broad interaction potential, some ZDHHC enzymes display striking selectivity, such as ZDHHC9. Although AlphaFold 3 predicts that GOLGA7B adopts a binding mode nearly identical to GOLGA7 in complex with ZDHHC9, our experimental data show that ZDHHC9 does not bind GOLGA7B. The structural basis for this selectivity is not apparent from current models, suggesting that specificity may be determined by subtle differences in surface features or conformational dynamics not captured by prediction alone. Unlike ZDHHC5, ZDHHC9 contains unique loop conformations and surface charge features near the interface region, which may contribute to differential recognition of GOLGA7 versus GOLGA7B. Key differences between Zdhhc5 and ZDHHC9 are observed at two variable domains: the N-terminus and the TM2-TM3 cytosolic loop. First, the N-terminus (residues 9-15) of Zdhhc5 appears flexible and is not modeled, whereas the expanded N-terminus in ZDHHC9 (residues 9-32) adopts a more complex loop-like structure. Second, the α-helix (α1) on the Zdhhc5 cytosolic loop between TM2 and TM3 was shorter than that on ZDHHC9, which formed a more distinctive loop of a different orientation. These variations between Zdhhc5 and ZDHHC9 were not structurally essential for interaction with GOLGA7 but may account for the inability of ZDHHC9 to interact with GOLGA7B. These differences were predicted in the AlphaFold 3 reference model and persisted in our cryo-EM model; however, the resolution in these regions was limited, and it is unclear whether they reflect true structural divergence or model variability. Higher-resolution data or additional validation will be necessary to determine their functional significance.

Our analysis revealed that deletion of any major portion of GOLGA7 disrupted complex formation with Zdhhc5. However, sequence alignment allowed us to identify the conserved RDYS motif as important for complex formation and stabilization of the zinc-finger domains, an insight supported by our Zdhhc5-GOLGA7 model. Notably, mutation of Pro^150^ in ZDHHC9, which directly contacts the GOLGA7 RDYS motif, is a cause of X-linked intellectual disorders (18, 31). This raises the possibility that other mutations disrupting RDYS-mediated binding will likewise result in similar neurodevelopmental disorders. More generally, our findings suggest that both the identity and *S*-acyation state of accessory proteins fine-tune ZDHHC binding specificity. Notably, we find that ZDHHC5, ZDHHC8, and ZDHHC9 each exhibit distinct interaction profiles with GOLGA7 and GOLGA7B, demonstrating that even among closely related PATs, interaction specificity is not conserved. Future studies of other ZDHHC enzymes interacting with GOLGA7 and GOLGA7B could uncover further regulatory complexity and specificity within this critical protein modification pathway and may reveal new therapeutic targets in diseases involving *S-*acylation.

Our findings also offer insight into how GOLGA7 regulates ZDHHC enzyme activity. In our cryo-EM structure, the GOLGA7 RDYS motif makes direct contact with the zinc finger motifs that stabilize the DHHC active site, suggesting a structural mechanism by which GOLGA7 may enhance catalytic function. Prior studies in yeast have shown that the Erf4 accessory protein, the GOLGA7 ortholog, enhances the catalytic activity of its DHHC partner Erf2 by slowing hydrolysis and thus stabilizing the active site intermediate (13). Similarly, GOLGA7 has been reported to be essential for ZDHHC9 palmitoylation activity (16). The fact that ZDHHC20, which lacks a requirement for an accessory protein, contains a membrane-inserted α5 helix in the region where GOLGA7 interacts with ZDHHC5 and ZDHHC9, further supports the notion that accessory proteins may functionally compensate for structural features absent in certain DHHC enzymes (9). Accessory proteins like GOLGA7 contribute to DHHC protein stability and membrane trafficking (13, 32), raising the possibility that GOLGA7 both promotes activity and protects ZDHHC5 from destabilization. Together, these observations reinforce the emerging paradigm in which accessory proteins dynamically tune the function, stability, and substrate engagement of membrane-modifying enzymes such as ZDHHC5.

## ACKNOWLEDGEMENTS

We thank L. Magtanong and L. Leak for comments. M.A.K. was supported in part by the NIH (T32GM007276). S.J.D. is supported by the NIH (R01CA272485).

## AUTHOR CONTRIBUTIONS

Conceptualization, M.A.K., J.A.B., and S.J.D.; Methodology, M.A.K., J.R., V.G., and H.W.; Investigation, M.A.K.; Writing – Original Draft, M.A.K. and S.J.D.; Statistics, D.B.; Writing – Review & Editing, J.R., L.L., V.G., and J.A.B.; Funding Acquisition and Supervision, J.A.B., and S.J.D.

## METHODS

### Cell lines and culture conditions

293T (sex: female) (ATCC CRL-3216), HEK 293S GnTI^-^ cells (sex: female) (ATCC CRL-3022), and Sf9 cells (ATCC CRL-1711) were obtained from ATCC (Manassas, VA) expanded for one passage, aliquoted, frozen at −80°C and stored in liquid nitrogen for subsequent experiments. HT-1080^N^ (sex: male) cells were described previously(33). HT-1080^N^ CRISPR/Cas9 Control, *ZDHHC5^KO1/2^* and *GOLGA7^KO1/2^*, and HEK 293T CRISPR/Cas9 Control, *ZDHHC5^KO1/2^* and *GOLGA7^KO1/^*^2^ cell lines were described previously (2). All HT-1080 cell lines were cultured in Dulbecco’s modified Eagle high-glucose plus pyruvate medium (DMEM, Cat# MT-10-013-CV, Gibco) with 10% (v/v) fetal bovine serum (FBS, Cat# 26140-079, Gibco), 0.5 U/mL Pen/Strep (P/S, Cat#5070-063, Gibco), and 1x non-essential amino acids (NEAAs, Cat# 11140-050, Gibco). All 293T cell lines were cultured in DMEM supplemented with 10% FBS (v/v) and 0.5 U/mL P/S. All HT-1080 and 293T cell lines were grown at 37°C with 5% CO_2_ in humidified tissue culture incubators (Thermo Fisher Scientific). All HT-1080 and 293T media were filtered through a 0.22 μM PES filter (Genesee Scientific) before use. Trypsin (Cat# 25200114, Gibco) was used for passaging of adherent cells. HT-1080 and 293T cells were counted using a Cellometer Auto T4 cell counter (Nexcelom, Lawrence, MA). HEK293S GnTI^-^ cells used for large-scale protein preps and CryoEM were grown in suspension at 37°C with 8% CO_2_ in Freestyle 293 medium (Cat# 12338018, Gibco) supplemented with 2% (v/v) FBS and 0.5 U/mL P/S. Sf9 cells used for baculovirus production and Sf9 Easy Titer (Sf9-ET) (ATCC CRL-3357) cells used for titering were grown in suspension at 27°C in Sf900 III SFM (Cat# 12658027, Gibco), and SF-X Insect Medium (Cat# SH30278.02, Hyclone) containing 50 μg/mL G418 (Cat# 10131-035, Gibco), respectively. HEK293S and Sf9 cells were counted manually.

### Chemicals and reagents

CIL56 was synthesized by Acme Bioscience (Palo Alto, CA). TOFA (Cat# T6575) was from Sigma-Aldrich (St. Louis, MI). SYTOX Green was from Life Technologies (Cat# S7020, Carlsbad, CA). All compounds were prepared as stock solutions in DMSO (Cat# 276855, Sigma-Aldrich) and stored at −20°C until use.

### Nuclear mKate2-Expressing Cell Lines

Cell lines stably expressing nuclear-localized mKate2 (denoted by the superscript “N”) were generated by lentiviral transduction. Previously established.(34) HEK 293T CRISPR/Cas9 Control cells and HEK 293T *GOLGA7* gene-disrupted (“KO”) cells were seeded at a density of 3 x 10^4^ cells per well in a twelve-well dish the day before transduction. Non-infected cells were also seeded separately for a simultaneous 2-fold puromycin dose response. 24 h later, the medium was replaced with 1 mL medium containing polybrene (8 µg/mL, Cat# H9268-5G, Sigma-Aldrich) and viral particles (Essen BioScience NLR puromycin red, Cat# 4625, Ann Arbor, MI) at a multiplicity of infection (MOI) of 3 TU/cell, calculated based on the manufacturer’s titer. 24 h later, the medium was removed and replaced with fresh medium. After 48 h, puromycin (10 µg/mL, Cat# A11138-03, Life Technologies) was added at a concentration determined by the puromycin dose response with non-infected cells (between 1-10 μg/mL). All non-transduced control cells were dead by day 3, and remaining infected cells were then transferred to larger plates in selection medium, expanded, and frozen down. HEK 293T CRISPR Control and HEK 293T *ZDHHC5^KO1/2^* were seeded and mKate2 stably expressing cell lines were generated with a bleomycin-selectable lentiviral construct (Cat# 4627, Essen BioScience). After infection with bleomycin-selectable lentivirus with the above protocol, cell lines were sorted to isolate mKate2-positive cells using a FACSAria II Fluorescence Activated Cell Sorter (BD Biosciences, San Jose, CA) at the Stanford Shared FACS Facility.

### Sequence alignment of PAT proteins

Clustal O v1.2.4 online(35) was used to generate a multiple sequence alignment of all mouse Zdhhc-family amino acid sequences using the following UniProtKB accession numbers: Q8R0N9 (Zdhhc1), P59267 (Zdhhc2), Q8R173 (Zdhhc3), Q9D6H5 (Zdhhc4), Q8VDZ4 (Zdhhc5), Q9CPV7 (Zdhhc6), Q91WU6 (Zdhhc7), Q5Y5T5 (Zdhhc8), P59268 (Zdhhc9), Q14AK4 (Zdhhc11), Q8VC90 (Zdhhc12), Q9CWU2 (Zdhhc13), Q8BQQ1 (Zdhhc14), Q8BGJ0 (Zdhhc15), Q9ESG8 (Zdhhc16), Q80TN5 (Zdhhc17), Q5Y5T2 (Zdhhc18), Q810M5 (Zdhhc19), Q5Y5T1 (Zdhhc20), Q9D270 (Zdhhc21), Q5Y5T3 (Zdhhc23), Q6IR37 (Zdhhc24), Q810M4 (Zdhhc25). Percent amino acid identity conservation between mouse and human PATs were determined by NCBI blastp (36) online multiple sequence alignment using the following accession numbers for the human enzymes: Q8WTX9 (ZDHHC1), Q9UIJ5 (ZDHHC2), Q9NYG2 (ZDHHC3), Q9NPG8 (ZDHHC4), Q9C0B5 (ZDHHC5), Q9H6R6 (ZDHHC6), Q9NXF8 (ZDHHC7), Q9ULC8 (ZDHHC8), Q9Y397 (ZDHHC9), Q9H8X9 (ZDHHC11), Q96GR4 (ZDHHC12), Q8IUH4 (ZDHHC13), Q8IZN3 (ZDHHC14), Q96MV8 (ZDHHC15), Q969W1 (ZDHHC16), Q8IUH5 (ZDHHC17), Q9NUE0 (ZDHHC18), Q8WVZ1 (ZDHHC19), Q5W0Z9 (ZDHHC20), Q8IVQ6 (ZDHHC21), Q8IYP9 (ZDHHC23), and Q6UX98 (ZDHHC24). Statistical tests were performed using the Mann–Whitney U test (two-tailed) using R v.4.4.3 through RStudio Desktop v.2024.12.1+563. To map conserved motifs unique to GOLGA7 binders, the aforementioned accession numbers were used to acquire text files of amino acid sequences for all 23 mouse Zdhhc proteins. Python v.3.12.8 was used to identify conserved sites grouped by length from these sequence files that were unique to GOLGA7 binders.

### Molecular biology

The following Zdhhc mouse enzyme expression vectors(21) were the kind gift of Dr. Masaki Fukata: pEF-BOS-HA, pEF-BOS-HA-DHHC1 (BC026570, Zdhhc1), pEF-BOS-HA-DHHC2 (NM_178395, Zdhhc2), pEF-BOS-HA-DHHC3 (NM_026917, Zdhhc3), pEF-BOS-HA-DHHC4 (NM_028379, Zdhhc4), pEF-BOS-HA-DHHC5 (NM_144887, Zdhhc5), pEF-BOS-HA-DHHC6 (NM_025883, Zdhhc6), pEF-BOS-HA-DHHC7 (NM_133967, Zdhhc7), pEF-BOS-HA-DHHC8 (AY668947, Zdhhc8), pEF-BOS-HA-DHHC9 (AK032233, Zdhhc9), pEF-BOS-HA-DHHC10 (AY668948, Zdhhc11), pEF-BOS-HA-DHHC11 (AY668949, Zdhhc23), pEF-BOS-HA-DHHC12 (BC021432, Zdhhc12), pEF-BOS-HA-DHHC13 (BC071194, Zdhhc24), pEF-BOS-HA-DHHC14 (BC059814, Zdhhc14), pEF-BOS-HA-DHHC15 (NM_175358, Zdhhc15), pEF-BOS-HA-DHHC16 (XM_129300, Q9ESG8, Zdhhc16), pEF-BOS-HA-DHHC17 (NM_172554, Zdhhc17), pEF-BOS-HA-DHHC18 (AY668950, Zdhhc18), pEF-BOS-HA-DHHC19 (BC049761, Zdhhc19), pEF-BOS-HA-DHHC20 (AY668951, Zdhhc20), pEF-BOS-HA-DHHC21 (NM_026647, Zdhhc21), pEF-BOS-HA-DHHC22 (NM_028031, Zdhhc13), and pEF-BOS-HA-DHHC23 (BC049767, Zdhhc25). pEF-BOS-HAZdhhc5^DHHS^, Zdhhc5^3CS^, and Zdhhc5^3YA^ expression vectors were described previously(2). pEF-BOS-3xHA-Zdhhc5 (described previously(2)), the Q5 site-directed mutagenesis kit (Cat# E0554, NEB, Ipswich, MA), and the *Z5 RNFR, Z5 ANAA, Z5 ANYR, Z5 ANAR, Z5 ANYA, Z5 RNAR, Z5 RNAA,* and *Z5 RNYA* primer sets (Table S1) were used to generate derivatives containing the following mutations in the Zdhhc5 coding sequence: Zdhhc5^RNFR^, Zdhhc5^ANAA^, Zdhhc5^ANYR^, Zdhhc5^ANAR^, Zdhhc5^ANYA^, Zdhhc5^RNAR^, Zdhhc5^RNAA^, and Zdhhc5^RNYA^ (targeting the R144-R147 ‘RNYR’ domain). pCI-neo-Flag-Zdhhc5 (37) obtained from Addgene (plasmid #85812, Watertown, MA), the NEB Q5 site-directed mutagenesis kit, and the *Z5ΔN-term*, *Z5ΔDHHC*, and *Z5ΔC-term* primer sets (Table S1) were used to generate derivatives containing the following mutations in the Zdhhc5 coding sequence: Δ14-270 (“Zdhhc5^ΔN-term^”), Δ104-154 (“Zdhhc5^ΔDHHC^”), and Δ270-714 (“Zdhhc5^ΔC-term^”). pEF-BOS-3xHA-DHHC4 and pEF-BOS-3xHA-DHHC19 plasmids, together with the Q5 site-directed mutagenesis kit and the *DHHC4 RNYR* and *DHHC19 RNYR* primer sets (Table S1), were used to generate Zdhhc4^RNYR^ and Zdhhc19^RNYR^ mutants that contained the RNYR motif and flanking residues. The Zdhhc4 mutant contained a substitution starting at A188, AWNTRYFLIY to RRNYRYFFLF, and the Zdhhc19 mutant starting at H151, HRNFRLFM to RRNYRYFF.

GOLGA7b-3xFLAG (Hs) was generated in the same backbone as our pDEST-pcDNA-GOLGA7-FLAG (Hs) construct. A gene block for use in Gibson assembly was designed from human GOLGA7b cDNA (NCBI Reference Sequence: NM_001010917.3) and ordered from Integrated DNA Technologies (IDT, Coralville, IA). The synthetized sequence contained a NotI site directly before the start codon for *GOLGA7b*, a TEV protease site immediately following the final *GOLGA7b* codon, and a three-residue linker (Gly, Arg, Ala) which contains an AscI site, a 3xFLAG tag, a stop codon, and a HindIII restriction site, allowing for removal of the entire cassette. The gene block was flanked by an AflII site which occurs 23 bp upstream of the *GOLGA7* start codon in the original vector, and XhoI site, which is immediately after the stop codon in the vector. For use as the backbone, pDEST-pcDNA5-GOLGA7-FLAG (described previously(2)) was digested with AflII (Cat# R0520, NEB) and Xho1 (Cat# R0146, NEB), followed by gel purification. NEB’s Gibson assembly kit (Cat# E5510S) was used to insert the *GOLGA7b* gene block into the empty vector and subsequently transformed in NEB Stable *E. coli* (Cat# C3040H). From this new GOLGA7b-3xFLAG plasmid, GOLGA7b^2CS^, with mutations Cys78Ser and Cys81Ser, and GOLGA7b^RDYS^ (R25-S28 to AAAA) were generated using the Q5 site-directed mutagenesis kit and the *GOLGA7b-C78,81S* and *GOLGA7b-RDYSΔAAAA* primer sets, respectively (Table S1).

pDEST-pCDNA-GOLGA7-3xFLAG was generated using the pDEST-pcDNA-GOLGA7b-3xFLAG backbone. The *GOLGA7* gene was PCR amplified from pDEST-pcDNA-GOLGA7-FLAG (ref.(2)) using *GOLGA7* Gibson insert primers (Table S1) with overlapping ends for Gibson assembly. An empty vector with complimentary ends for Gibson assembly was generated by PCR amplification of pDEST-pcDNA-GOLGA7b-3xFLAG using *GOLGA7 Gibson vector primers* (Table S1) and inserting the *GOLGA7* PCR fragment using NEB’s Gibson assembly kit (Cat# E5510S).

pcDNA-GOLGA7^2CS^-3xFLAG construct containing a TEV protease site and three residue linker (Gly, Arg, Ala) connecting the C-terminus GOLGA7^2CS^ to the 3x-FLAG tag *was generated using the GOLGA7^2CS^-3xFLAG* primer pair (Table S1), the previously described(2) GOLGA7^2CS^-FLAG construct, and the Q5 site-directed mutagenesis kit (Cat# E0554, NEB). Four GOLGA7 deletion mutants (GOLGA7^Δ10-35^, GOLGA7^Δ35-65^, GOLGA7^ΔC-term^, GOLGA7^ΔN-term^) were generated from pDEST-pcDNA5-GOLGA7-FLAG (ref.(2)) using the Q5 site-directed mutagenesis kit and the *G7d10-35, G7d35-65, G7dC-term*, and *G7dN-term* primer pairs (Table S1). These primer pairs deleted segments Gly10-Glu35, Glu35-Tyr65, K91-R137, and P3-L30 from GOLGA7, respectively.

To generate N-St2-sfGFP-PPX-Zdhhc5 and N-FLAG-GOLGA7 constructs for CryoEM, *GOLGA7* (Hs) and Z*dhhc5* (Mm) were cloned from pEF-BOS-3xHA-Zdhhc5 and pDEST-pcDNA5-GOLGA7-FLAG plasmids (described previously)(2), into a pEG BacMam vector as described(38) to allow for baculovirus generation and large-scale expression. First, the Q5 site-directed mutagenesis kit (Cat# E0554, NEB) was used to add AscI and NotI restriction sites flanking each gene, allowing for easy transfer to BacMam plasmids. This was done using the *Z5 AscI/Z5 NotI*, and *G7 AscI/G7 NotI* primer sets, respectively (Table S1). Next, restriction digests of these two plasmids were performed with Asc I (Cat# FD1894, Thermo Fisher) and Not I (Cat# FD0593, Thermo Fisher) to obtain *GOLGA7* and *Zdhhc5* inserts. These inserts were gel purified and *GOLGA7* was ligated into an AscI/NotI digested and the gel purified N-FLAG BacMam vector generated and described previously (39) containing an N-terminal FLAG tag. *Zdhhc5* was ligated into an AscI/NotI digested and gel purified N-St2-sfGFP BacMam vector generation and described previously (39), which contained an N-terminal Strep II tag, superfolder GFP (40), and an HRV 3C protease site preceding the gene. These ligations were performed according to manufacturer protocol using Thermo Scientific T4 DNA Ligase (Cat# EL0016). All DNA gel purifications were performed using 1% agarose gel and the Qiagen QIAquick Gel Extraction Kit (Cat# 28704).

### Transfection, cell lysis and coimmunoprecipitation

For coimmunoprecipitation (co-IP) studies, 500,000 HEK 293T cells were seeded in 6-well plates 24 h prior to transfection, with 2 wells per construct. Each well containing 1 mL of media was transfected with 2.5 μg plasmid DNA, 6 μL Lipofectamine LTX Reagent, and 2.5 µL PLUS Reagent (Cat# 15338030, Invitrogen) in 250 μL Opti-MEM Reduced Serum Media (Cat# 31985-062, Life Technologies) according to manufacturer’s instructions, incubated overnight, and transfection media replaced with 2 mL fresh media the following day. After 24 h in fresh media, cells were washed once with HBSS (Cat# 14025-134, Life Technologies) and resuspended in 1 mL HBSS. Each pair of wells was combined and pelleted (2348 x *g*, 1 min), and the supernatant removed and discarded. Cell pellets were stored at −20°C until lysis. Pellets were resuspended in lysis buffer (50 mM Tris-HCl pH 7.4, 150 mM NaCl, 2 mM EDTA, P8340 Protease Inhibitor Cocktail EDTA-free (100x) (Cat# 11836170001, Roche, Basel, Switzerland), 1% n-dodecyl-β-maltoside (DDM, Cat# D310, Anatrace, Maumee, OH)), and passed through a 25-gauge needle 20 times to homogenize. Lysates were spun (18,000 x *g*, 15 min, 4°C), supernatant removed and the pellet discarded. Protein levels were quantified using the Pierce BCA protein assay (Cat# 23228 and 23224, Thermo Fisher). For pulldowns, 90 μg of cell lysate was set aside for SDS gel input. Lysate was added to 15 μL pre-washed and equilibrated Anti-FLAG(R) M2 magnetic beads (Cat# M8823, Sigma-Aldrich, 50% slurry) or anti-HA magnetic beads (Cat# 88837, Pierce, Thermo Fisher) at 1 mg/mL in 900 mL and incubated at 4°C overnight with end-over-end rotation. Beads were washed three times with 1 mL TBS (20 mM Tris base, 150 mM NaCl, pH 7.4) containing 0.5% n-dodecyl-β-maltoside before elution into 30 μL Bolt LDS Sample Buffer (Cat# B0007, Life Technologies). Elution was carried out by vortexing for 5 min at 4°C followed by incubation for 30 min at 37°C. Eluted proteins were analyzed by immunoblotting.

### Immunoblotting

90 µg of total protein (input) was combined with 4x Bolt LDS Sample Buffer, incubated in a 37°C water bath for 30 min, and loaded onto a Bolt 4-10% Bis-Tris Plus Gel (Cat# NW04120BOX, Life Technologies). Eluates from the co-IP were loaded alongside the respective inputs. The Chameleon Duo Ladder (Cat# 928-60000, LI-COR, Lincoln, NE) was used as a standard. Gels were run at 100 V for 1.5 h or until the dye front reached the end, and proteins were transferred to nitrocellulose membrane using the iBlot2 transfer stack (Cat# IB23002, Invitrogen). Membranes were blocked with Odyssey Blocking Buffer (Cat# 927-50000, LI-COR) (30 min, RT), and incubated in primary antibody mixture (overnight, 4°C), except for GOLGA7 primary antibody (1 h RT). Membranes were washed three times in Tris buffered saline (20 mM Tris base, 150 mM NaCL, pH 7.4) with 0.1% Tween 20 (TBST), then incubated with secondary antibody mixture (1 h, RT). Following secondary incubation, membranes were washed 3 times in TBST and then scanned on a two-channel Odyssey CLx Imaging System (LI-COR). All antibodies were diluted in Odyssey Blocking Buffer. Antibodies used were rabbit α-ZDHHC5 (1:250, Cat# HPA014670, Sigma-Aldrich), mouse α-GOLGA7 (1:250, Cat# H00051125, Novus Biologicals, Littleton, CO), rabbit α-ZDHHC9 (1:250, Cat# NBP1-84499, Novus Biologicals), rabbit α-FLAG (1:1000, Cat# 14793S, Cell Signaling Technology), rabbit α-HA (1:1000, Cat# 3724S, Cell Signaling Technology, Danvers, MA) and mouse α-C4 actin (1:4000, Cat# SC47778, Santa Cruz Biotechnology), donkey α-rabbit (1:15,000, Cat# 926-32213, 926-68023, LI-COR), donkey α-goat (1:15,000, Cat# 926-32214, 926-68024, LI-COR), and donkey α-mouse (1:15,000, Cat# 926-32212, 926-68022, LI-COR).

### GOLGA7 binding heatmap calculation

Using Image J, protein densitometry was calculated from the mouse Zdhhc enzyme co-IP screen for endogenous GOLGA7. Immunoblots were quantified for GOLGA7 input and pulldown, Zdhhc enzyme input and pulldown, and background. After pixel inversion and background subtraction for each band, percentage of total GOLGA7 recovered by each enzyme was calculated by dividing each GOLGA7 pulldown band by its respective input band. Percent of each enzyme occupied by GOLGA7 was also calculated by dividing GOLGA7 pulldown (G7^PD^) by Zdhhc pulldown (Zdhhc^PD^). This was normalized to Zdhhc5/GOLGA7 binding by dividing percent enzyme occupied by GOLGA7 (G7^PD^/Zdhhc^PD^) by percent Zdhhhc5 occupied (G7^PD^/Zdhhc5^PD^) as follows:

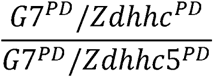

Morpheus (https://software.broadinstitute.org/morpheus) was used to generate a heatmap representing percent GOLGA7 binding for each mammalian PAT enzyme relative to Zdhhc5 binding.

### Cell death assessment using microscopy

Cell death was assayed using scalable time-lapse analysis of cell death kinetics (STACK) (33, 41). HEK 293T^N^ cell lines were seeded at 25,000 cells per well in 12 well plates. The next day, the medium was removed and replaced with medium containing 20 nM SYTOX Green (SG, Cat# S7020, Life Technologies) ± lethal compounds and inhibitors. Cells were imaged at 4 h intervals for 96 h using the Essen IncuCyte ZOOM live-cell analysis system (Essen BioSciences). Counts of mKate2-positive objects (mKate2^+^, live cells) and SG-positive objects (SG^+^, dead cells) were counted using IncuCyte ZOOM Live-Cell Analysis System software (Essen Biosciences). Image analysis parameter values were as follows for mKate2^+^ objects: parameter adaptive, threshold adjustment 1; Edge split on; Edge sensitivity 50; Filter area min 20 mm^2^, maximum 8,100 mm^2^; Eccentricity max 1.0; and SG^+^ objects: Parameter adaptive, threshold adjustment 10; Edge split on; Edge sensitivity −5; Filter area min 20 mm^2^, maximum 750 mm^2^; Eccentricity max 0.9. Lethal fraction scores were computed in Microsoft Excel from mKate2^+^ and SG^+^ counts as described (33, 41). For any given timepoint *n* in a treatment time course (t = 0 ➔ t = n) consisting of mKate2^+^, SG^+^, and double positive (SG^+^/mKate2^+^) cells, the lethal fraction (LF) was calculated as described (41) and by Equation 1.

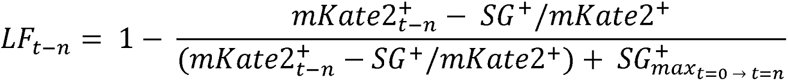

For HEK 293T *Zdhhc5* and *Golga7* gene-disrupted cell lines that did not express mKate2, only SG^+^ objects were counted. The following image extraction parameters were used: Parameter adaptive, threshold adjustment 1; Edge split on; Edge sensitivity −10; Filter area min 5 mm^2^, maximum 800 mm^2^; Eccentricity max 0.9.

### Immunofluorescence and confocal imaging

60,000 293T *ZDHHC5^KO^* cells were seeded per well in a 12-well plate on #1.5 glass coverslips coated with poly-D30 lysine 24 h prior to transfection. To each well containing 1 mL of medium was added 1 μg plasmid DNA, 3 μL Lipofectamine 3000 Reagent, and 6 µL P3000 Reagent (Cat# L3000001, Invitrogen) in 125 μL Opti-MEM Reduced Serum Media (Cat# 31985-062, Life Technologies) according to manufacturer’s instructions, incubated overnight, and transfection media replaced with 1 mL fresh media the following day. After 48 h in fresh media, coverslips were rinsed in 1x PBS and fixed in 4% paraformaldehyde (Alfa Aesar, Ward Hill, MA) for 20 min at room temperature. Cells were rinsed once with 1x PBS and permeabilized overnight at 4°C in 1x PBS with 3% BSA (Gemini Bio Products, West Sacramento, CA) and 0.1% Triton X-100 (“PBS-BT”). Coverslips were transferred to a hydration chamber and covered in 30 μL PBS-BT. 30 μL primary antibody mix was applied to coverslips for 1 h at room temperature. Primary antibodies used were rabbit α-ZDHHC5 (Cat# HPA014670, Sigma-Aldrich, 1:100) and mouse α-pan-cadherin (CH-19, ab6528, Abcam, Fremont, CA, 1:100). Following primary incubation, coverslips were washed 3 times with PBS-BT and incubated with 30 μL of secondary antibody mix (1:1000 of each secondary antibody, obtained from Life Technologies) for 1 h at room temperature. Secondary antibodies used were goat α-rabbit 568 (Cat# A11036, Life Technologies) and goat α-mouse 488 (Cat# A11029, Life Technologies). Coverslips were washed 5 times with PBS then mounted in ProLong Glass antifade reagent containing NucBlue stain (Cat# P36981, Thermo Fisher). Coverslips were cured overnight at room temperature in the dark before storage at 4°C and imaging. Confocal images were obtained on a Zeiss LSM780 laser scanning confocal microscope (Carl Zeiss) with a 63x/1.4 NA plan apochromat objective.

Zdhhc5 and pan-cadherin expression was robust across every condition except the Zdhhc5 negative control which displayed no signal. Fluorescence imaging was performed with identical acquisition settings across conditions, including fixed laser power, gain, and exposure time. Because different fluorescent channels exhibited varying signal intensities, only threshold levels were adjusted for each channel to optimize visualization while maintaining the dynamic range, with no adjustment to brightness or contrast. These adjustments ensured that signal was neither saturated nor undetectable. As such, adjustments were not and should not be used for quantitative comparisons between channels. Fluorescence intensity line profile analysis was performed using FIJI v2.16.0 (42) for three cells per condition across two replicates, but signal was used to determine spatial localization and colocalization only, not to compare relative expression across conditions or between channels.

### Homology modeling and structural analysis

AlphaFold 3 (25) was used to generate a model of mouse Zdhhc5 and human GOLGA7 in complex using amino acid sequences from the following UniProt accession numbers: Q8VDZ4 (Zdhhc5) and Q7Z5G4 (GOLGA7). The same was done for human ZDHHC5 (Q9C0B5) and GOLGA7. The ZDHHC9/GOLGA7 and Erf2/Erf4 CryoEM structures (18) were imported into ChimeraX v1.8 (43) with the following PDB accession numbers: 8HF3 and 8HFC. Structural comparison and figure generation were carried out in ChimeraX and PyMOL (The PyMOL Molecular Graphics System, Version 3.0 Schrödinger, LLC). Molecular graphics and analyses performed with ChimeraX.

### Protein expression and purification for CryoEM

P3 baculovirus containing N-St2-sfGFP-PPX-Zdhhc5 (Mm) and N-FLAG-GOLGA7 (Hs) BacMam constructs were generated in Sf9 cells as described (38) and used to infect HEK293S GnTI^-^ cells. HEK293S GnTI^-^ cells were grown in suspension until they reached a density of 3 × 10^6^ cells/ml and then infected with P3 baculovirus at a multiplicity of infection (MOI) of 1 for Zdhhc5 and 4 for GOLGA7. After 12 h, 10 mM sodium butyrate (Cat# B5887, Sigma-Aldrich) was added to the medium and the temperature reduced to 30°C. The cells were harvested 48 h later by centrifugation (4800 x *g*, 20 min), supernatant discarded, and pellets washed once in PBS (pH 7.5; Cat# 10010023, Gibco), and pelleted again (4800 x *g*, 20 min). Supernatant was discarded and cell pellets were frozen in liquid nitrogen and stored at −80°C until needed.

Cell pellets were thawed on ice and resuspended in 15 mL of lysis buffer per gram of cells. Lysis buffer was composed of 50 mM Tris/NaOH (pH 7.5), 375 mM NaCl, 1 μg/mL leupeptin (Cat# E18, Sigma-Aldrich), 1 μg/mL aprotinin (Cat# A6279, Sigma-Aldrich), 1 μg/mL pepstatin A (Cat# P5318, Sigma-Aldrich), 1 mM phenylmethylsulfonyl fluoride (Cat# 10837091001, Roche), 100x cOmplete EDTA-free protease inhibitor cocktail (Cat# 73567200, Roche), 5 mM β-mercaptoethanol (β-Me), 10% glycerol (w/v), and 10 μg/mL DNaseI (Cat# 04536282001, Roche). The homogenate was sonicated at 70% for 45 s (5 s on, 5 s off) with a probe sonicator, then the complex extracted by adding 1% (w/v) DDM with 0.2% (w/v) cholesterol hemisuccinate (CHS; Cat# C6512, Sigma-Aldrich) rotating for 2 h at 4°C. The mixture was clarified by centrifugation at 80,000 x *g* for 30 min, and the supernatant was added to 15 μL MagStrep StrepTactin Type III beads (Cat# 15362006, IBA Lifesciences, Germany) per gram of cells and rotated at 4°C for 1 h. The beads were collected on a magnetic rack, washed 3x with 5 column volumes (cv.) of 20 mM HEPES/HCl (pH 7.5), 150 mM NaCl (HBS) with 0.01% (w/v) glyco-diosgenin (GDN, Cat# 850525P, Avanti, Alabaster, AL). After removing wash buffer, 25 µL of the GDN/TBS buffer was added to the beads along with HRV 3C protease (Cat# 71493, Novagen, Madison, WI) at 10 U/mg of complex, and the complex was left to cut off the beads rotating overnight at 4°C. The cut complex was removed from the beads using a magnetic rack and hard spun at 20,000 x *g* for 20 min. Zdhhc5/GOLGA7 concentration was calculated from its absorbance at 280 nm assuming an extinction coefficient (ε280) of 57190 M^−1^ cm^−1^ (calculated by ProtParam33) (44). The sample was then concentrated to 2 mg/mL in a centrifugal tube (Amicon Ultra-4; 100-kDa cutoff, Millipore Sigma, Burlington, MA) and frozen on grids.

Protein quality was assayed by size-exclusion chromatography (SEC) and SDS-PAGE. For SEC, 4 μL of sample was injected onto a Superose 6 Increase column (Cytiva, Marlborough, MA) previously equilibrated with HBS with 0.01% (w/v) GDN and UV absorbance measured. For SDS-PAGE, 3 μL of cut final product and 10 μL of prep fractions collected (crude, flow-through, wash) were combined with 4x Bolt LDS Sample Buffer and 10% βMe, incubated in a 37°C water bath for 30 min, and loaded onto a Bolt 4-10% Bis-Tris Plus Gel. The Chameleon Duo Ladder was used as a standard. The gel was Coomassie stained overnight with SimplyBlue SafeStain (Cat# LC6060, Invitrogen), destained in water, and viewed on a BioRad ChemiDoc imager.

### Cryo-EM sample preparation and data acquisition

Cryo-EM grids were frozen using a Vitrobot Mark IV (FEI) as follows: 3 μL of the sample at RT was applied to a glow-discharged Quantifoil R1.2/1.3 holey carbon 400 mesh gold grid (Quantifoil Micro Tools, Germany), blotted for 2.5 - 4 s in 100% humidity at 22°C, and plunge frozen in liquid ethane cooled by liquid nitrogen. Grids were screened for ice thickness and particle distribution using a Glacios (200 kV; Thermo Scientific) in the Yale School of Medicine Center for Cellular and Molecular Imaging. Cryo-EM data were recorded on a Titan Krios (FEI) operated at 300 kV in the Yale West Campus Cryo-EM Core, equipped with a Gatan K2 Summit camera and imaging filter. SerialEM (version 4.1-beta) (45) was used for automated data collection. Movies were collected at a nominal magnification of 81,000× in super-resolution mode resulting in a calibrated pixel size of 0.5 Å per pixel, with a defocus range of approximately –0.8 to –2.5 μm. Fifty frames were recorded over 10 s of exposure at a dose rate of 1.67 electrons per square Å per frame. Video frames were aligned and binned over 2 x 2 pixels using MotionCor2 (46) and the contrast transfer function parameters for each motion-corrected image were estimated using CTFFIND4 (47).

### Cryo-EM data analysis and model building

For data processing, a flow chart can be found in **Supplementary Fig. 6.** In brief, 17,391 dose-weighted micrographs with 0.8 Å per pixel size were applied for patch-based CTF estimation, motion correction (48), auto-particle picking, and 2D classification in CryoSPARC (49) (Structural Biotechnology Inc.), yielding 1,383,157 particles with a box size of 384 pixels, binned over 2 x 2 pixels. After removal of junk particles, the remaining 694,239 particles were re-extracted without binning and subjected to three rounds of ab initio reconstruction and a non-uniform refinement (50), resulting in a 5.9 Å map, and finally local refinement yielding a 3.9 Å map. Using the AlphaFold 3 (25) model of Mm Zdhhc5 and Hs GOLGA7 in complex, the predicted structures were docked into the density map and manually adjusted in Coot v.0.9.8.8 (51). Images of the model were created using UCSF ChimeraX and PyMOL.

### Data availability

The final cryo-EM map has been deposited into the Electron Microscopy Data Bank under the accession number EMD-70274. The final model has been deposited into the PDB under accession number 9OA6. Coordinated for ZDHHC9/GOLGA7 and Erf2/Erf4, used for structural comparisons in this paper, were obtained from the PDB under accession numbers 8HF3 and 8HFC, respectively. For all other data requests, contact S.J.D.

### Software

Data were analyzed and graphed using Excel v.16.95.4 (Microsoft Corp.), GraphPad Prism v.10.4.0 (GraphPad Software Inc.), and Adobe Illustrator (Adobe Inc. v.29.4).

