## Supplemental Table 1 for "Functional Dissection of the Zdhhc5-GOLGA7 Protein Palmitoylation Complex"

Table S1 – Primers

| Oligo Name | Sequence | Source |
| --- | --- | --- |
| Z5 RNFR Fwd | ttcagaTACTTCTTCCTTTTCCTCC | This paper |
| Z5 RNFR Rev | gttcctGCGACCAATACAGTTGTTTAC | This paper |
| Z5 ANAA Fwd | gccgcaTACTTCTTCCTTTTCCTCC | This paper |
| Z5 ANAA Rev | gttggcGCGACCAATACAGTTGTTTAC | This paper |
| Z5 ANYR Fwd | TATTGGTCGCgccAACTACAGATACTTC | This paper |
| Z5 ANYR Rev | CAGTTGTTTACCCAAGGG | This paper |
| Z5 ANAR Fwd | gccagaTACTTCTTCCTTTTCCTCC | This paper |
| Z5 ANAR Rev | gttggcGCGACCAATACAGTTGTTTAC | This paper |
| Z5 ANYA Fwd | tacgcaTACTTCTTCCTTTTCCTCC | This paper |
| Z5 ANYA Rev | gttggcGCGACCAATACAGTTGTTTAC | This paper |
| Z5 RNAR Fwd | gccagaTACTTCTTCCTTTTCCTCC | This paper |
| Z5 RNAR Rev | gttcctGCGACCAATACAGTTGTTTAC | This paper |
| Z5 RNAA Fwd | gccgcaTACTTCTTCCTTTTCCTCC | This paper |
| Z5 RNAA Rev | gttcctGCGACCAATACAGTTGTTTAC | This paper |
| Z5 RNYA Fwd | tacgcaTACTTCTTCCTTTTCCTCC | This paper |
| Z5 RNYA Rev | gttcctGCGACCAATACAGTTGTTTAC | This paper |
| Z5ΔN-term Fwd | CCAGAAGTGTCAGATGGG | This paper |
| Z5ΔN-term Rev | ATACTTGCTGGGTTTGAAC | This paper |
| Z5ΔDHHC Fwd | TCCCTGACAGCTCACATC | This paper |
| Z5ΔDHHC Rev | TTTCATTCGCACCTGGATAC | This paper |
| Z5ΔC-term Fwd | TGAGCGGCCGCTTCCCTT | This paper |
| Z5ΔC-term Rev | TCGAAGGAAAGGAGGTCTGATTACAATTG | This paper |
| DHHC4 RNYR Fwd | tacttcttccttttcCTCCTGACGCTGACGGCT | This paper |
| DHHC4 RNYR Rev | cctgtagttcctgcgCCCGATGCAGTTGTTCACC | This paper |
| DHHC19 RNYR Fwd | cgctacttcttcCTGCTGGTCCTGTCACTC | This paper |
| DHHC19 RNYR Rev | gtagttgcggcgACCGATGCAGTTATTGACC | This paper |
| GOLGA7b-C78,81S Fwd | gccagcGCCACGGCCTACTTCATC | This paper |
| GOLGA7b-C78,81S Rev | caggctGCCCTCGAGGTAGGAGCT | This paper |
| GOLGA7b-RDYSDAAAA Fwd | gccgccGATGGGACCATCTGTCAG | This paper |
| GOLGA7b-RDYSDAAAA Rev | ggctgcCTGGATAAAGACCTTGGTG | This paper |
| G7 Gibson Ins Fwd | GCGGCCGCGCCACCatgaggccgcagcagg | This paper |
| G7 Gibson Ins Rev | CCCGCCCTGGAAGTAAAGGTTTTCtcttccactgctcatgcc | This paper |
| G7 Gibson Vec Fwd | GGTGGCGCGGCCGCtTAAGTTTAAACGC | This paper |
| G7 Gibson Vec Rev | GAAAACCTTTACTTCCAGGGCGGGCG | This paper |
| G7-2CS-3xFLAG Fwd | gatgacgatgacaaggactataaggatgatgacgataaaGATTACAAGGATGACGATGAC | This paper |
| G7-2CS-3xFLAG Rev | cttgtaatcggcgcgcccgccctggaagtaaaggttttcTCTTCCACTGCTCATGCC | This paper |
| G7d10-35 Fwd | AACCGGATTGATAGGCAGCAGTTTGAAGAAAC | This paper |
| G7d10-35 Rev | TCCGGACACCGGCGCCTG | This paper |
| G7d35-65 Fwd | CTCGAAGGTTGTTTGGCTTGTTTAAC | This paper |
| G7d35-65 Rev | CTCCAGCTCCGCAGGGAA | This paper |
| G7dC-term Fwd | TACCCAACTTTCTTGTAC | This paper |
| G7dC-term Rev | CAGAACCTTCTCATAATGAG | This paper |
| G7dN-term Fwd | GGCGGCCAGTCATATCTC | This paper |
| G7dN-term Rev | CCTCATGCCAACTTTTTTGTAC | This paper |
| Z5 AscI Fwd | gcgcCATGCCCGCAGAGTCTGG | This paper |
| Z5 AscI Rev | gccaTCCAGCGTAATCTGGAACATC | This paper |
| Z5 NotI Fwd | cgcctGAGGATCCGAATTAGAGTG | This paper |
| Z5 NotI Rev | gccgccACAGAAATCTCATAAGTGG | This paper |
| G7 AscI Fwd | ggcgcgCCATGAGGCCGCAGCAGG | This paper |
| G7 AscI Rev | TGGCGCGG**G**CGCTTAAGT | This paper |
| G7 NotI Fwd | cgccGAAAACCTTTACTTCCAGGGcggg**g**g | This paper |
| G7 NotI Rev | gccgcTCTTCCACTGCTCATGCC | This paper |

| Sequencing Primers |  |
| --- | --- |
| Z5_648 | cgaccagaagtttcagatgggc |
| Z5_1043 | cttggaaccagagagcttccg |
| Z5_1798 | ccttgcagccattactgacacc |
| EF-1aforward | TCAAGCCTCAGACAGTGGTTC |
