## Supplemental Figure 1 for "Functional Dissection of the Zdhhc5-GOLGA7 Protein Palmitoylation Complex"

**A**

Protein sequence conservation between mouse and human DHHC enzymes

| Zdhhc | 1 | 2 | 3 | 4 | 5 | 6 | 7 | 8 | 9 | 11 | 12 | 13 | 14 | 15 | 16 | 17 | 18 | 19 | 20 | 21 | 23 | 24 | 25 |
|---|---|---|---|---|---|---|---|---|---|---|---|---|---|---|---|---|---|---|---|---|---|---|---|
| % identity | 87 | 95 | 97 | 74 | 98 | 93 | 95 | 92 | 98 | 60 | 87 | 91 | 95 | 98 | 95 | 99 | 93 | 69 | 88 | 98 | 90 | 88 | NA |

**B**

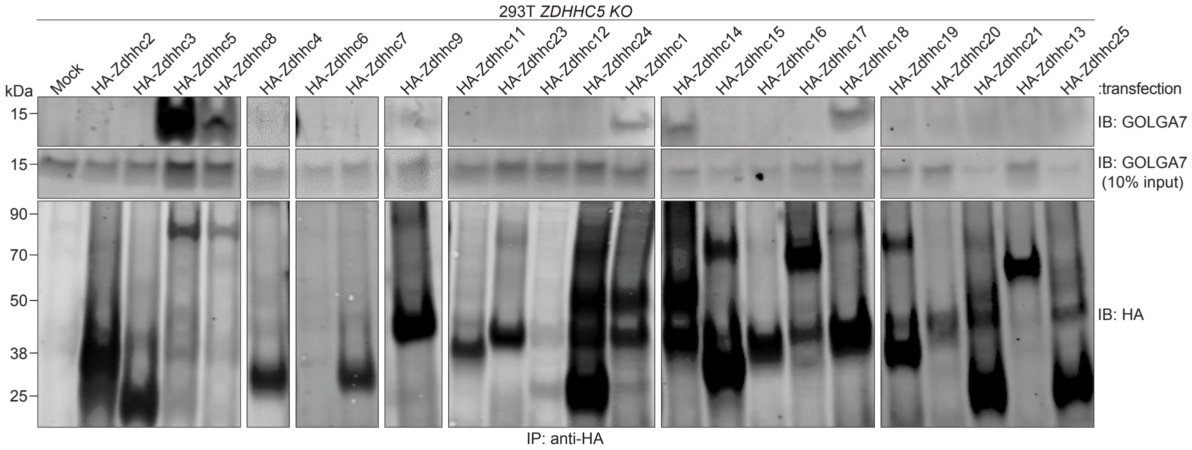


**C**

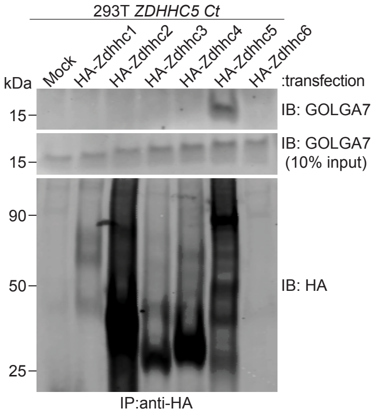


**Figure S1. Interaction between Zdhhc-family proteins and endogenous GOLGA7, related to Figure 1.** (A) Sequence conservation (% identity) between mouse Zdhhc enzymes and their human homologs. Mann-Whitney test, not significant (n.s.). (B) Immunoblot analysis from co-immunoprecipitation samples transfected as indicated. These blots are representative of two to three independent experiments and are from a large-scale screen. Due to the number of conditions, samples were transfected and processed in parallel across multiple experiments. Representative lanes were compiled from blots run and developed under identical conditions. Some samples are shown as intact blots, while others appear as single or double cropped lanes; this reflects differences in sample order and the presence of replicates on the same blot. Cropped lanes are delineated by white lines. All shown data derive from the same experimental series and were processed in parallel. Note: data for Zdhhc14-18 are also shown in Main Figure 1. (C) Immunoblot analysis from co-immunoprecipitation samples transfected as indicated. Blot is representative of two independent experiments.
