## Supplemental Figure 2 for "Functional Dissection of the Zdhhc5-GOLGA7 Protein Palmitoylation Complex"

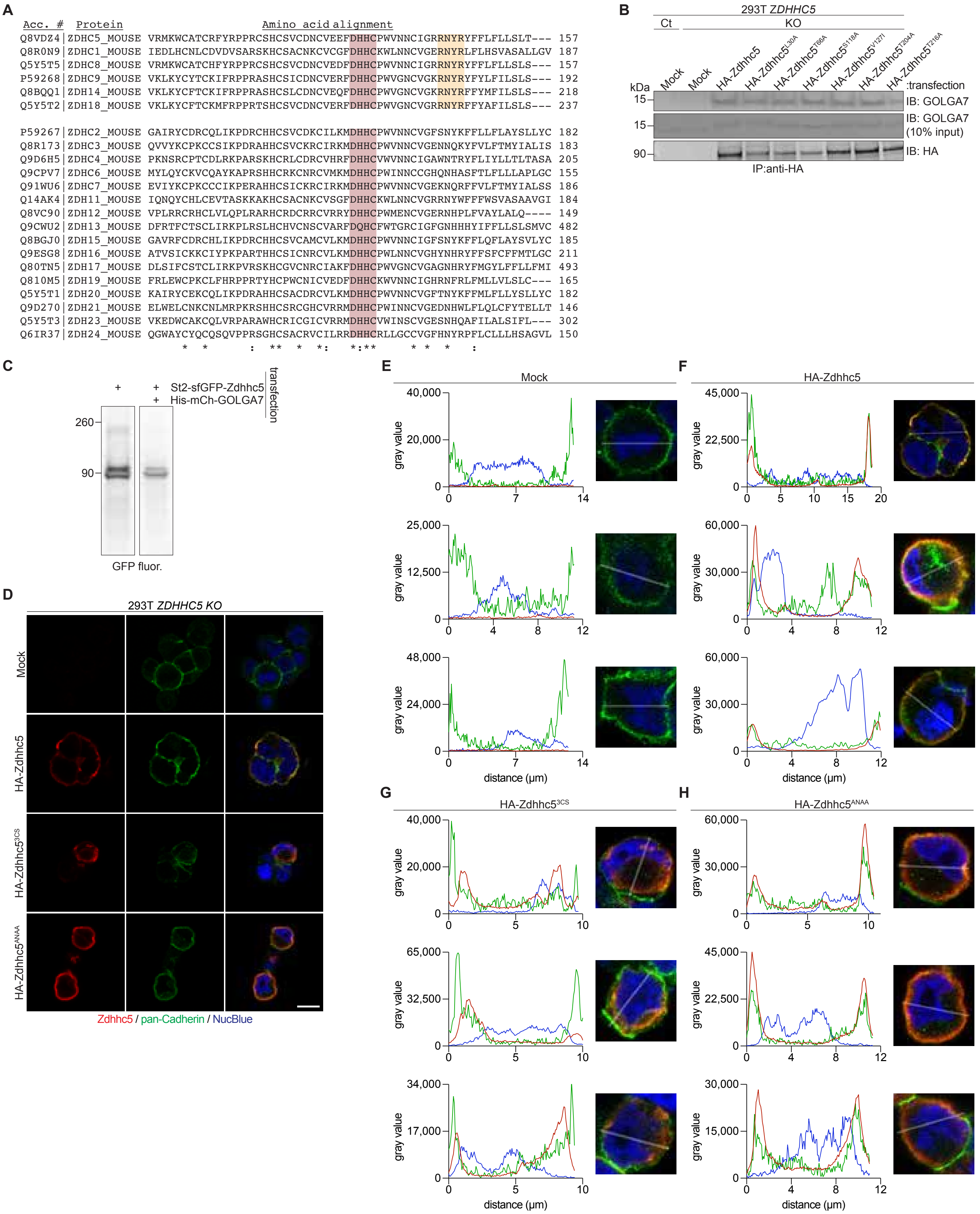


**Figure S2. Interaction between Zdhhc5 mutants and endogenous GOLGA7, related to Figure 2.** (A) Clustal Omega amino acid alignment of Zdhhc family conserved cysteine rich domain and DHHC active site (red), highlighting the RNYR motif in conserved GOLGA7 binders (top) vs. nonbinders (bottom). (B) Immunoblot analysis of co-immmunoprecipitation samples in Control (Ct) and *Zdhhc5* gene disrupted ("KO") cells transfected as indicated. Representative of two independent experiments. (C) GFP fluorescence of lysate from 293T cells transfected as indicated. Lanes shown side-by-side were cropped from the same blot and are shown adjacent for clarity. Representative of two independent experiments. (D) Immunofluorescence of 293T *Zdhhc5 KO* cells transfected as indicated. (E-H) Fluorescence intensity line profile analysis of cells corresponding to (D). Three cells were quantified per condition. (D-H) Red=Zdhhc5, Green=pan-Cadherin, Blue=NucBlue. Representative of two independent experiments.
