## Supplemental Figure 3 for "Functional Dissection of the Zdhhc5-GOLGA7 Protein Palmitoylation Complex"

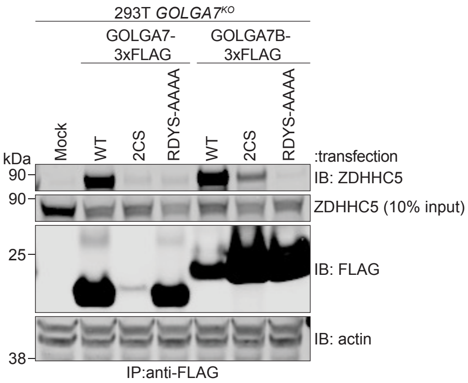


**Figure S3. ZDHHC5 interaction with GOLGA7 and GOLGA7B mutants, Related to Figure 3.** Co-immuno-precipitation analysis of ZDHHC5 binding in *GOLGA7* gene disrupted (”KO”) cell lines transfected as indicated.
