## Supplemental Figure 4 for "Functional Dissection of the Zdhhc5-GOLGA7 Protein Palmitoylation Complex"

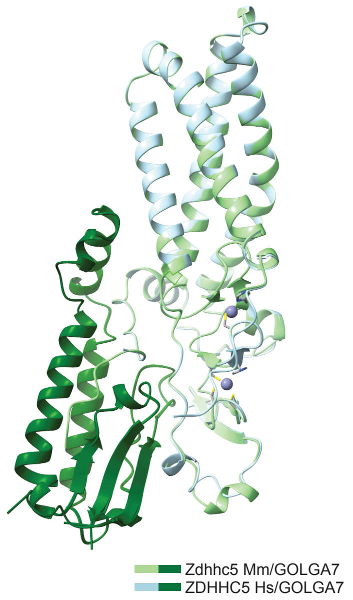


**Figure S4. AlphaFold3 models of human and mouse ZDHHC5/GOLGA7 complexes, related to Figure 5.** AlphaFold3 generated model of *H. sapiens* (Hs) ZDHHC5 and *M. musculus* (Mm) Zdhhc5 in complex with Hs GOLGA7, generated using the following UniProt accessions: Q9C0B5 (Hs ZDHHC5), Q8VDZ4 (Mm Zdhhc5), Q7Z5G4 (Hs GOLGA7).
