## Supplemental Figure 5 for "Functional Dissection of the Zdhhc5-GOLGA7 Protein Palmitoylation Complex"

```
enzyme                     amino acid alignment
ZDHHC9  23   FCCDGRVMMARQKGIFYLTLFL-ILGTCTLFFAFECRYLAVQLS--PAIPVFAAMLFLFS  79
             F   GR    +     +L + L I+    LF  FE   L    +    + +F    ++ +
Erf  2  61   FFLGGRFRTVKGAKPLWLGVLLAIVCPMVLFSIFEAHKLWHTQNGYKVLVIFFYYFWVIT  120

ZDHHC9  80   MATLLRTSFSDPGVIPRALPDEAAFIEMEIEATNGAVPQGQRPPPRIKNFQINNQIVKLK  139
             +A+ +RT+ SDPGV+PR        I +     N  +PQ       +      ++ + +K
Erf  2  121  LASFIRTATSDPGVLPRN-------IHLSQLRNNYQIPQEYYNLITLPTHSSISKDITIK  173

ZDHHC9  140  YCYTCKIFRPPRASHCSICDNCVERFDHHCPWVGNCVGKRNYRYFYLFILSLSLLTIYVF  199
             YC +C+I+RPPR+SHCS C+ CV   DHHC WV NC+GKRNYR+F +F+L   L ++ +
Erf  2  174  YCPSCRIWRPPRSSHCSTCNVCVMVHDHHCIWVNNCIGKRNYRFFLIFLLGAILSSVILL  233

ZDHHC9  200  AFNIVYVALKSLKIGFLETLKETPGTVLEVLICFFTLWSVVGLTGFHTFLVALNQTTNED  259
                 +++A +S         ++ P  +L +     TLW    L  +H F+    QTT E
Erf  2  234  TNCAIHIARES------GGPRDCPVAILLLCYAGLTLWYPAILFTYHIFMAGNQQTTREF  287

ZDHHC9  260  IKGSWTGKNRV-------QNPYSHGNIVKNCCEVLCGPLPPSVLDRR  299
             +KG  + KN V       +N Y+ G+ +KN   ++  P  PS +  R
Erf  2  288  LKGIGSKKNPVFHRVVKEENIYNKGSFLKNMGHLMLEPRGPSFVSAR  334
```

**Figure S5. Amino acid alignment.** Clustal Omega amino acid alignment of human ZDHHC9 and yeast ERF2. Red highlights the conserved DHHC active site; orange highlights the RNYR site conserved among GOLGA7 binders.
