## Supplemental Figure 6 for "Functional Dissection of the Zdhhc5-GOLGA7 Protein Palmitoylation Complex"

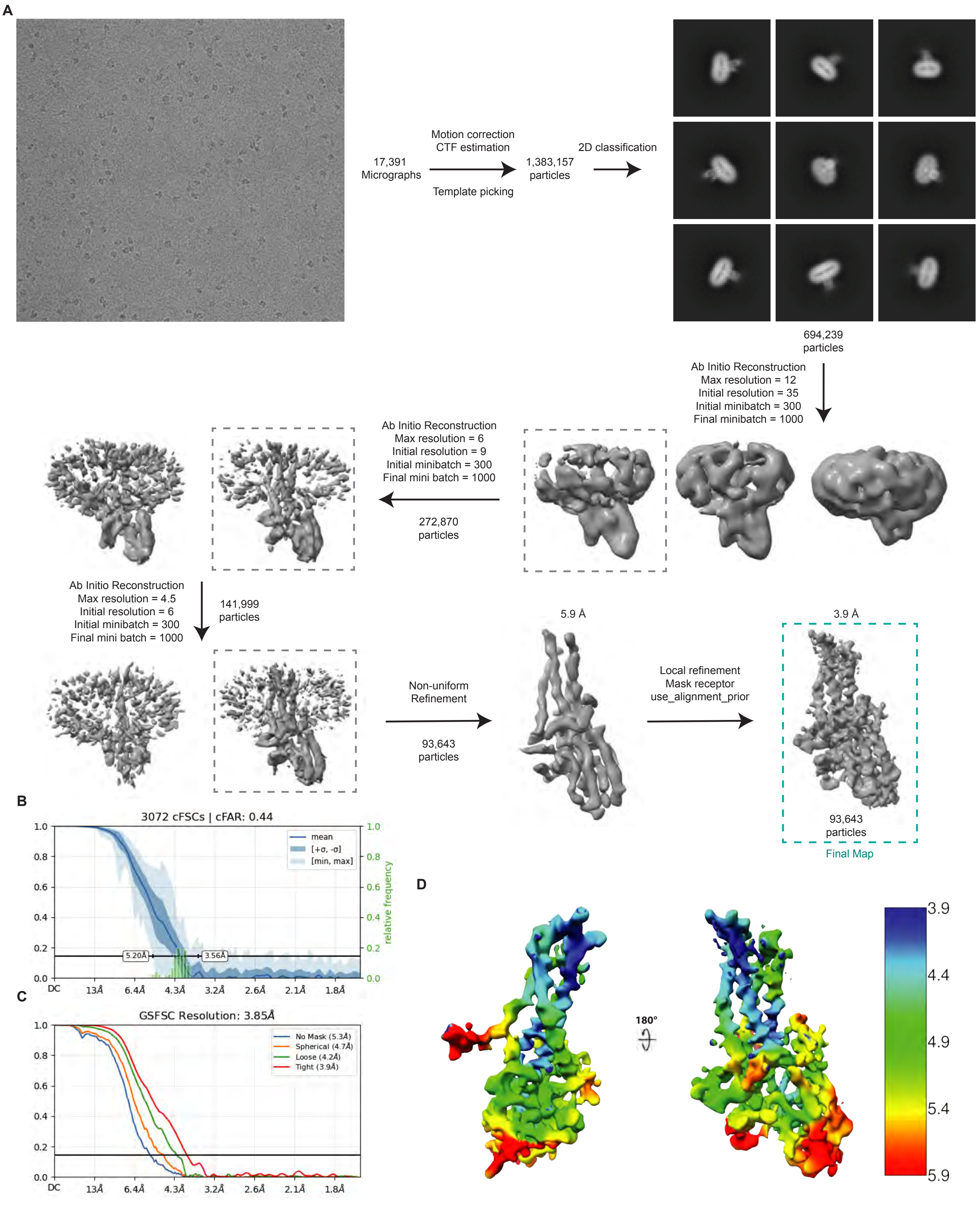


**Figure S6. Cryo-EM data analysis of the Zdhhc5-GOLGA7 complex.** (A) A representative cryo-micrograph from the chosen grid and flowchart of EM data processing including representative two-dimensional class averages. (B) Directional corrected FSC (cFSC) curves from CryoSPARC showing resolution anisotropy across 3D orientations. A cFAR value of 0.44 indicates moderate anisotropy. Relative frequency (right y-axis) reflects orientation sampling density. (C) Gold standard Fourier shell correlation (GSFSC) curves for final 3D reconstruction. FSC curves were calculated in cryoSPARC using different masking conditions: no mask (blue), spherical mask (orange), loose mask (green), and tight mask (red). The reported resolution (3.85 Å) is defined at the 0.143 FSC threshold.
